# FACT is recruited to the +1 nucleosome of transcribed genes and spreads in a Chd1-dependent manner

**DOI:** 10.1101/2020.08.20.259960

**Authors:** Célia Jeronimo, Andrew Angel, Christian Poitras, Pierre Collin, Jane Mellor, François Robert

## Abstract

The histone chaperone FACT occupies transcribed regions where it plays prominent roles in maintaining chromatin integrity and preserving epigenetic information. How it is targeted to transcribed regions, however, remains unclear. Proposed models for how FACT finds its way to transcriptionally active chromatin include docking on the RNA polymerase II (RNAPII) C-terminal domain (CTD), recruitment by elongation factors, recognition of modified histone tails and binding partially disassembled nucleosomes. Here, we systematically tested these and other scenarios in *Saccharomyces cerevisiae* and found that FACT binds transcribed chromatin, not RNAPII. Through a combination of experimental and mathematical modeling evidence, we propose that FACT recognizes the +1 nucleosome, as it is partially unwrapped by the engaging RNAPII, and spreads to downstream nucleosomes aided by the chromatin remodeler Chd1. Our work clarifies how FACT interacts with genes, suggests a processive mechanism for FACT function, and provides a framework to further dissect the molecular mechanisms of transcription-coupled histone chaperoning.

**Highlights:**

- High-resolution mapping of FACT localization in yeast
- FACT binds partially unwrapped nucleosomes in transcribed genes, not RNAPII
- FACT distribution along genes requires Chd1
- Processive mechanism for FACT function

## Introduction

Nucleosomes, the building blocks of chromatin, allow for DNA compaction and the regulation of protein-DNA interactions. Nucleosomes also represent roadblocks to molecular machines that translocate along DNA. For instance, during transcription elongation, RNA polymerase II (RNAPII) encounters successions of nucleosomes, which must unfold for the DNA template to feed the polymerase active site. Importantly, however, these nucleosomes have to reassemble in the wake of the polymerase to maintain chromatin integrity (Lai and Pugh, 2017; Teves et al., 2014; Venkatesh and Workman, 2015). In yeast, this delicate process is regulated by several chromatin regulators including histone methyltransferases (Set2), histone demethylases (Rpd3S), chromatin remodelers (Chd1 and others) and histone chaperones (FACT, Spt6 and others) (Lai and Pugh, 2017; Venkatesh and Workman, 2015). Among those, histone chaperones FACT and Spt6 play master roles during transcription elongation as they can mediate both nucleosome disassembly and assembly by their capacity to bind both DNA and histones.

To assist RNAPII during transcription elongation through nucleosomes, chromatin regulators such as histone chaperones need to be recruited to sites of transcription. Most of these factors (Spt6, Set2, and Rpd3S) are recruited to these sites via their interaction with RNAPII, with important contributions from its phosphorylated C-terminal domain (CTD) (Carrozza et al., 2005; Drouin et al., 2010; Govind et al., 2010; Krogan et al., 2003; Sdano et al., 2017). How FACT makes its way to actively transcribed regions, however, remains unclear. An interaction with RNAPII or an RNAPII-associated protein would provide a simple way to recruit the histone chaperone to the right locations. Accordingly, several proteins have been proposed to recruit FACT to transcribing RNAPII, including the phosphorylated RNAPII CTD (Kwon et al., 2010; Mason and Struhl, 2003), the elongation factor PAF1 complex (PAF1C) (Adelman et al., 2006), the capping enzyme Cet1 (Sen et al., 2017), the SetD2 methyltransferase (Carvalho et al., 2013), the histone acetyltransferase NuA3 (John et al., 2000) and the chromatin remodeler Chd1 (Krogan et al., 2002), but the relative contribution of these factors to FACT recruitment has never been systematically analyzed. Alternatively, FACT may recognize transcribed chromatin rather than the elongation complex itself. Such a model would explain why FACT is recruited to transcription sites by RNAPI and III in addition to RNAPII (Birch et al., 2009; Tessarz et al., 2014). Recruitment through transcribed chromatin is supported by the recent analysis of the DNA fragments associated with FACT after micrococcal nuclease (MNase) treatment (Martin et al., 2018). These data, however, do not rule out the possibility of a transient interaction with the polymerase prior to more stable (or more easily crosslinked) interactions with nucleosomes. Also, the biochemical nature of such a “transcription-modified nucleosome” that would be recognized by FACT remains to be determined. Previous studies suggested that histone acetylation, methylation or ubiquitylation may contribute to FACT recruitment (Carvalho et al., 2013; Fleming et al., 2008; Pathak et al., 2018), potentially providing a molecular mechanism for how transcription –by co-transcriptional histone modifications– would promote the recruitment of FACT to transcribed chromatin. Alternatively, *in vitro* studies showing increased affinity of FACT for disrupted nucleosomes (Formosa et al., 2001; Liu et al., 2020; McCullough et al., 2018; Nesher et al., 2018; Ruone et al., 2003; Tsunaka et al., 2016; Wang et al., 2018) suggest that partial loss of histone-DNA contacts, generated by RNAPII or some associated factors, may allow FACT binding.

Here, we systematically tested many possible recruitment mechanisms, including those proposed previously, and concluded that FACT binds transcribed chromatin rather than RNAPII or elongation factors. We, however, found no evidence for a role of histone modifications in FACT recruitment. Instead, our data support a model where partial unwrapping of the first nucleosome (+1 nucleosome), induced by engaging RNAPII, exposes a FACT interacting surface on the nucleosome and triggers its recruitment. Once docked on the +1 nucleosome, FACT appears to spread to more downstream nucleosomes with the help of the chromatin remodeler Chd1, suggesting that FACT works processively along genes during their transcription.

## Results

### FACT is recruited after transcription initiation in an RNAPII CTD phosphorylation-independent manner

To start investigating how FACT is recruited to chromatin, we compared the distribution of FACT to that of RNAPII and nucleosomes at high-resolution. We performed chromatin immunoprecipitation (ChIP), followed by high-resolution tiling arrays (ChIP-chip) and sequencing exonuclease-treated ChIP (ChIP-exo) of FACT subunits (Spt16 and Pob3) and RNAPII (Rpb3) in wild type (WT) cells (**Figures 1** and **S1**). In agreement with previous work (Martin et al., 2018; Mason and Struhl, 2003; Pathak et al., 2018; Vinayachandran et al., 2018), we found both Spt16 and Pob3 to occupy the transcribed region of transcriptionally active genes (**Figures 1A** and **S1A**). Both FACT subunits correlated with RNAPII occupancy (**Figures 1A, 1B and S1B**). Moreover, shutting down transcription using the *rpb1-1* allele led to a nearly complete loss of FACT occupancy genome-wide (**Figure 1C**). A comparison of the data for RNAPII and Spt16 and Pob3 indicates that FACT accumulates downstream of RNAPII (**Figures 1A** and **S1A**). Indeed, FACT occupancy peaks at each nucleosome dyad within transcribed regions while RNAPII occupancy builds up upstream of the +1 nucleosome and has a more uniform distribution further downstream (**Figure 1A**, metagenes on the left). Thus, genome-wide FACT occupancy correlates with − and depends on− transcription, but does not precisely correlate with where transcription initiates. This observation, which corroborates ChIP-exo, ChIP-seq, and MNase-ChIP-seq data by others (Martin et al., 2018; Vinayachandran et al., 2018), suggests that FACT “joins” RNAPII post-initiation.

**Figure 1:**
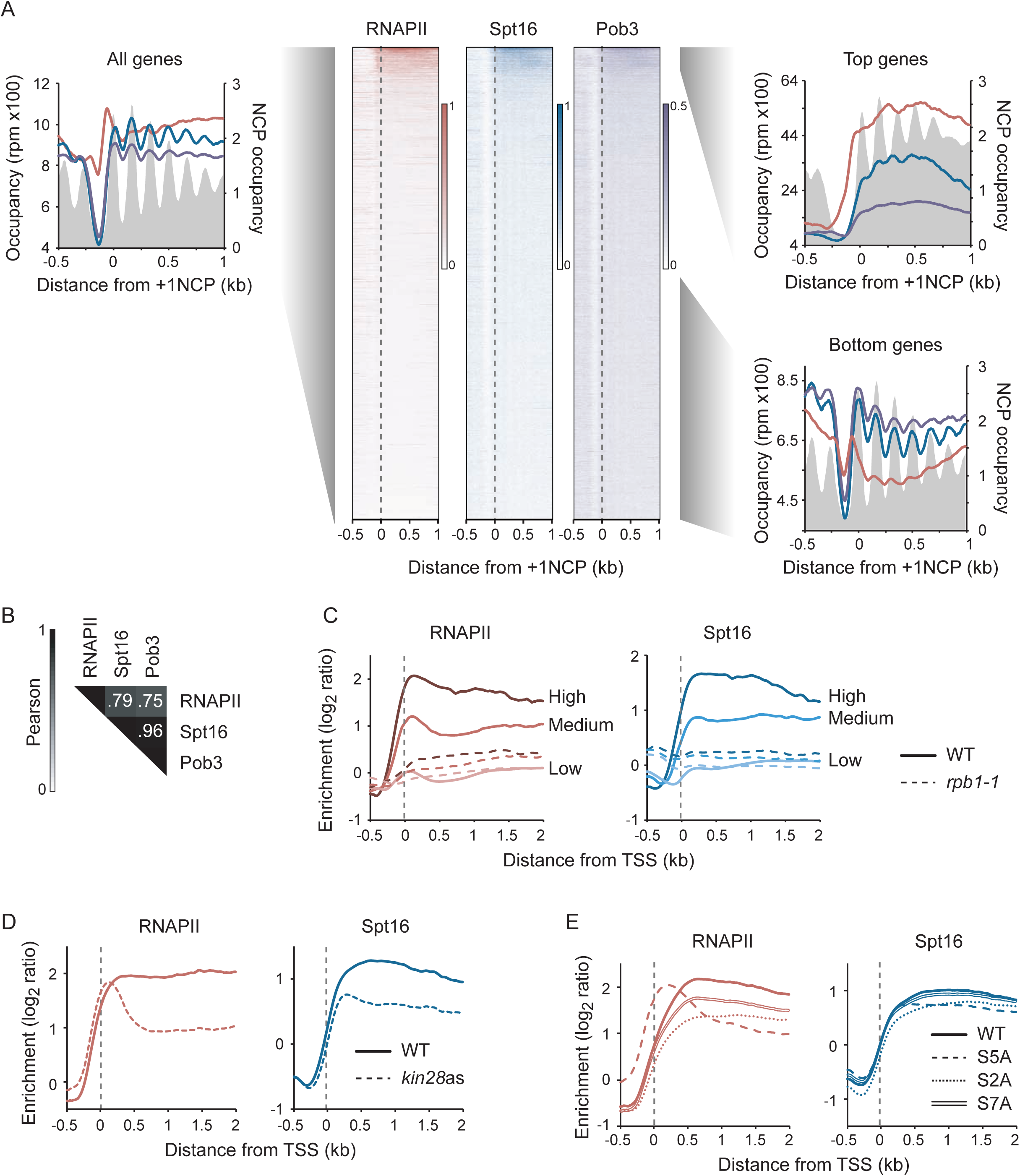
FACT occupies actively transcribed regions and is recruited after initiation in a CTD-phosphorylation independent manner. **A**) Metagene and heatmap representations of RNAPII (Rpb3) and both FACT subunits (Spt16, Pob3) occupancy, as determined by ChIP-exo, in WT cells. Metagenes for all genes (n = 5,456) as well as the most transcribed genes (Top genes; n = 264) and least transcribed (Bottom genes; n = 2,792) are shown. The average enrichment level of nucleosomes (Jiang and Pugh, 2009) is shown in grey. Genes in the heat maps were ordered by decreasing RNAPII occupancy (n = 5,456). Data are aligned on the dyad position of +1 nucleosomes, as determined in (Chereji et al., 2018). **B**) Pearson correlation matrix of RNAPII (Rpb3), Spt16 and Pob3 occupancy, as determined by ChIP-exo. Pearson correlations were calculated using all of the data points covering nuclear ORFs that are longer than 1KB (n = 3503). **C**) Metagenes of RNAPII (Rpb3,) and FACT (Spt16,) occupancy over highly (dark shade; n = 85), mildly (medium shade; n = 190) and lowly (light shade; n = 3,241) transcribed genes, as determined by ChIP-chip, relative to Input, in WT (solid traces) and *rpb1-1* (dashed traces) cells after an 80 min heat-shock at 37°C. **D**) Metagenes of RNAPII (Rpb3) and FACT (Spt16) occupancy over transcribed genes (n = 275), as determined by ChIP-chip, relative to Input, in WT (solid traces) and *kin28* ATP analog-sensitive (*kin28*as, dashed traces) cells, both treated 15 min with NAPP1. **E**) Metagenes of RNAPII (Rpb3) and FACT (Spt16) occupancy over transcribed genes (n = 355), as determined by ChIP-chip, relative to Input, in cells ectopically expressing the indicated CTD versions of *RPB1* (WT, solid traces; S5A, dashed traces; S2A, dotted traces; S7A thin traces) following nuclear depletion of the endogenous Rpb1 protein by anchor-away (after a 90 min treatment with rapamycin, see STAR Methods). Data for RNAPII in panels D and E (except for the S2A mutant) are from (Jeronimo and Robert, 2014). NCP, nucleosome core particle. TSS, transcription start site. See also **Figure S1**.

Complete pre-initiation complexes (PICs) are too transient to be detected by ChIP in WT cells (Jeronimo and Robert, 2014; Wong et al., 2014) so that FACT may be a transient component of PICs that escapes detection in WT cells. To test this possibility, we performed RNAPII and FACT ChIP-chip experiments under two conditions that stabilize PICs: inhibition of TFIIH kinase (Kin28) and mutation of serine 5 of the RNAPII CTD. Inhibition of Kin28 and the CTD-S5A mutation both led to the accumulation of RNAPII in the promoter region as expected (Jeronimo and Robert, 2014; Wong et al., 2014) (**Figures 1D** and **1E**). In contrast, while FACT occupancy is decreased on gene bodies (reflecting reduced RNAPII occupancy) it does not shift upstream (**Figures 1D** and **1E**). These results do not support a direct role for FACT in transcription initiation genome-wide, although we cannot exclude roles at specific genes.

FACT has been proposed to be recruited to the elongating RNAPII via CTD phosphorylation by TFIIH (Mason and Struhl, 2003). The finding that Spt16 occupancy is unaffected by either Kin28 inhibition or CTD-S5A mutations (except for effects explained by changes in RNAPII occupancy) (**Figures 1D** and **1E)**, does not support such a model. Of note, we also observed no FACT occupancy defects in cells expressing CTD-S2A or CTD-S7A mutants (**Figure 1E**). Finally, we profiled FACT occupancy in various CTD kinase mutants including *ctk1Δ, cdk8Δ, bur2Δ* and the double *kin28*as/*bur2Δ* and found no effect on FACT occupancy that could not be solely explained by an effect on RNAPII occupancy (**Figure S1C**). We, therefore, conclude that FACT is unlikely to be recruited to genes via the phosphorylated CTD.

### FACT occupancy is uncoupled from RNAPII in *CHD1* mutants

Some evidence suggested that FACT may be recruited to the elongating RNAPII via interactions with the elongation factor PAF1C (Adelman et al., 2006), or the N-terminal domain of the capping enzyme Cet1 (Sen et al., 2017). We, therefore, tested FACT (Spt16) and RNAPII (Rpb3) occupancy by ChIP-chip in PAF1C mutants (*paf1Δ, cdc73Δ, rtf1Δ, ctr9Δ*, and *leo1Δ*), and in cells where the N-terminal domain of Cet1 is truncated (Cet1ΔN). None of these mutations alter FACT localization (**Figures S2A and S2B)**. We conclude that neither the PAF1C nor the N-terminal domain of Cet1 are required to target FACT to transcribed regions.

The data presented above are consistent with FACT occupancy on transcribed genes simply reflecting the presence of transcription, without a requirement for a recruiting factor. But what makes transcribed chromatin recognized by FACT? One possibility is that transcription generates a specific histone modification pattern that is then recognized by FACT. Several studies provided evidence for such a scenario (Fleming et al., 2008; John et al., 2000; Pathak et al., 2018; Smart et al., 2009). We therefore profiled FACT (Spt16) and RNAPII (Rpb3) occupancy by ChIP-chip in *sas3Δ* (the catalytic subunit of NuA3), *gcn5Δ, set2Δ, bre1Δ* (the E3 ligase for H2B ubiquitylation), *ubp8Δ* (the main deubiquitylase for H2B) and *ubp10Δ* (a FACT-dependent H2B deubiquitylase) cells. As shown in **Figure S2C**, however, none of these mutants altered FACT occupancy. We conclude that histone modifications do not contribute significantly to FACT recruitment during transcription, although we can not rule out the possibility that a modification we have not tested may contribute.

Previous work showed that FACT physically interacts with the ATP-dependent chromatin remodeler Chd1 (Farnung et al., 2017; Krogan et al., 2006; Krogan et al., 2002; Krogan et al., 2004; Simic et al., 2003). To investigate the possible role of Chd1 in FACT recruitment, we performed ChIP-chip and ChIP-exo of FACT in *chd1Δ* cells (**Figures 2A** and **S2D**). Surprisingly, the deletion of *CHD1* had a profound effect on FACT occupancy. In *chd1Δ* cells, FACT occupancy increased in the 5’ end of genes (**Figures 2A** and **S2D**). These data were observed in two different genetic backgrounds (W303 and BY4741; **Figure S2D**) and were not observed in mutants for the two other yeast chromatin remodelers involved in nucleosome sliding, Isw1 and Isw2 (**Figure S2E**). Importantly, this change in FACT occupancy occurred without a change in RNAPII occupancy (**Figures 2A** and **S2D)**. Hence, FACT occupancy is uncoupled from RNAPII occupancy in *chd1Δ* cells, providing direct evidence that FACT interacts with chromatin rather than with RNAPII. This conclusion is further supported by profiles of FACT occupancy in *spt6-1004*, a mutant of the Spt6 histone chaperone that affects histone occupancy in the 5’ end of genes more so than in the 3’ end (Jeronimo et al., 2019; Jeronimo et al., 2015). Indeed, in *spt6-1004* cells, while RNAPII occupancy is reduced uniformly along genes, Spt16 and Pob3 show a biased decrease in the 5’ end of genes, mirroring the occupancy defect of histones in this mutant (**Figure S2F**).

**Figure 2:**
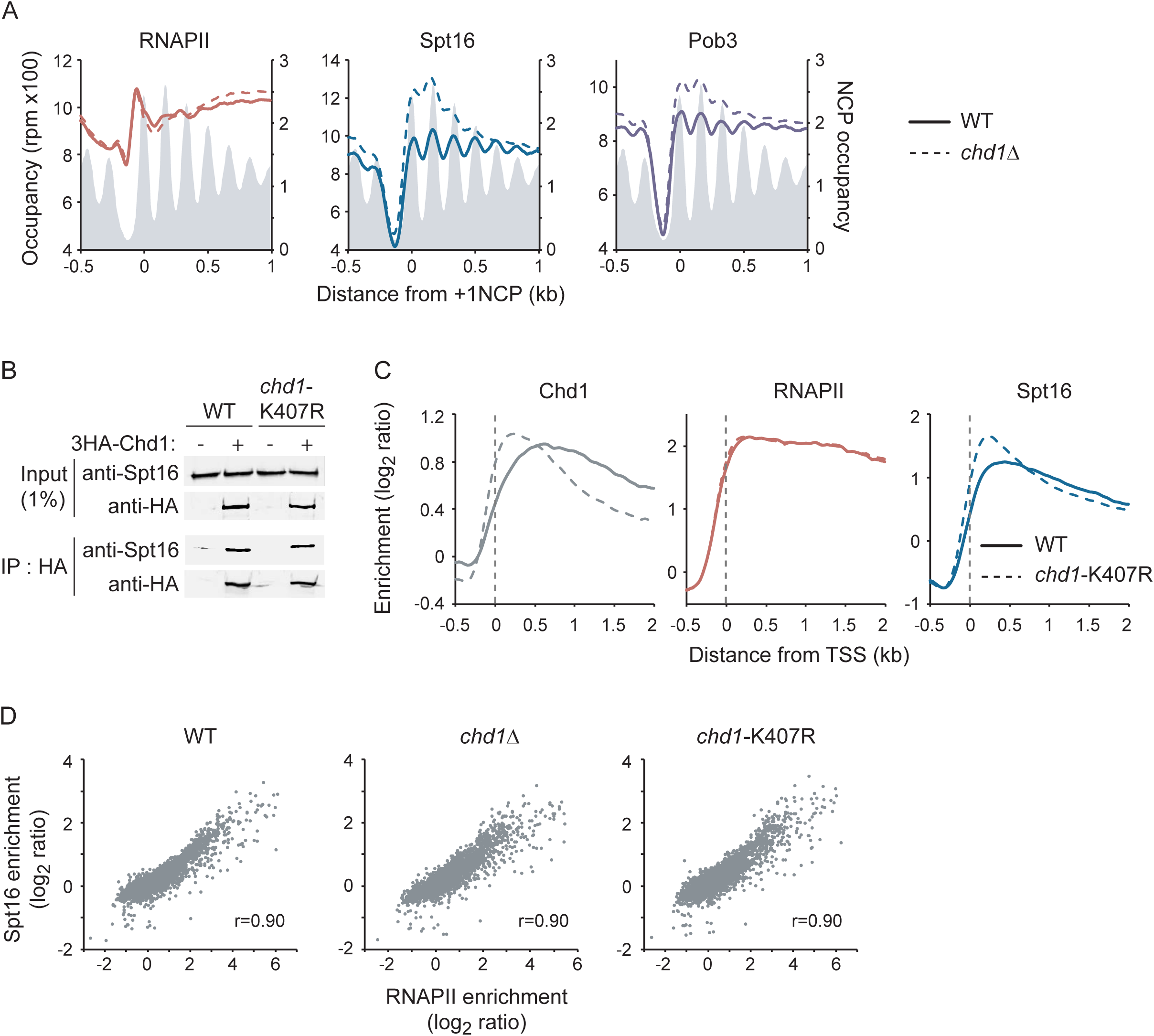
FACT, but not RNAPII, accumulates at 5’ nucleosomes in CHD1 mutants. **A**) Metagene representation of RNAPII (Rpb3) and FACT (Spt16, Pob3) occupancy over genes (n = 5,456), as determined by ChIP-exo, in WT (solid traces) and *chd1*Δ (dashed traces) cells. The average enrichment level of nucleosomes (Jiang and Pugh, 2009) is shown in grey. Data are aligned on the dyad position of +1 nucleosomes (Chereji et al., 2018). **B**) Co-immunoprecipitation experiments of FACT (Spt16) and Chd1 in WT and *chd1-*K407R cells expressing an HA-tagged version of Chd1. **C**) Metagene representation of Chd1, RNAPII (Rpb3), and FACT (Spt16) occupancy over transcribed genes (n = 275), as determined by ChIP-chip, relative to Input, in WT (solid traces) and *chd1*-K407R (dashed traces) cells. **D**) Scatter plots of the ORFs occupancy (n = 5817) of FACT (Spt16) versus RNAPII (Rpb3) in WT, *chd1*Δ, and *chd1*-K407R cells. NCP, nucleosome core particle. TSS, transcription start site. See also **Figure S2**.

To determine whether Chd1 affects FACT occupancy through protein-protein interactions or through its effect on chromatin, we tested a mutant in the helicase/ATPase domain (*chd1*-K407R) that is predicted to prevent Chd1 from remodeling nucleosomes (Simic et al., 2003). This mutant does not affect the physical interaction of Chd1 with FACT (**Figure 2B**) but phenocopies the effect of *chd1Δ* on FACT occupancy (**Figure 2C**). This result strongly suggests that Chd1 impacts FACT occupancy via its effect on chromatin rather than by directly recruiting the histone chaperone. We noted, however, that even in *chd1Δ* and *chd1*-K407R cells, FACT occupancy correlated with transcription (**Figure 2D**), demonstrating that FACT recruitment is transcription-dependent even in that context.

The accumulation of FACT at the 5’ end of genes when Chd1 or its remodeling activity is disrupted is inconsistent with the model that FACT simply recognizes and binds to disrupted nucleosomes wherever they occur. Instead, it suggests an unexpected mechanism in which FACT is recruited to the 5’ end of genes in an RNAPII-independent but transcription-dependent manner, and spreads through the gene body via a mechanism that requires the remodeling activity of Chd1. This mechanism is explored below.

### FACT binds disorganized nucleosomal particles on transcribed genes

Because all our data point to recognition of altered nucleosomes produced by transcription, and, perhaps, by Chd1, as key to determining where FACT binds, we investigated the structure of the nucleosomes bound by FACT *in vivo*. We performed MNase-ChIP-seq of FACT (Spt16 and Pob3) in both WT and *chd1*Δ cells. MNase digestion mainly generates fragments of nucleosome size (∼150 base pairs (bp)) but previous work showed that insights about non-canonical nucleosomal particles, such as partially unwrapped nucleosomes, can be obtained by analyzing DNA fragments deviating from 150 bp (Henikoff et al., 2011; Martin et al., 2018; Ramachandran et al., 2017). We therefore scrutinized the size of the mapped DNA fragments from the FACT (Spt16 and Pob3) MNase-ChIP-seq experiments. As expected, the Input material contains mainly mononucleosomal-size DNA, with some dinucleosomes (**Figures 3A**, grey shades). Interestingly, however, the Spt16 ChIPs in both WT and *chd1*Δ cells were enriched for DNA fragments of subnucleosomal (<150 bp) and internucleosomal (150-300 bp) sizes (**Figure 3A**, blue traces). The enrichment of these DNA fragments relative to mononucleosomes is even more pronounced when looking only at fragments aligning on the most highly transcribed genes (**Figure 3A**, right panel). Similar results were obtained with Pob3 (**Figure S3A**).

**Figure 3:**
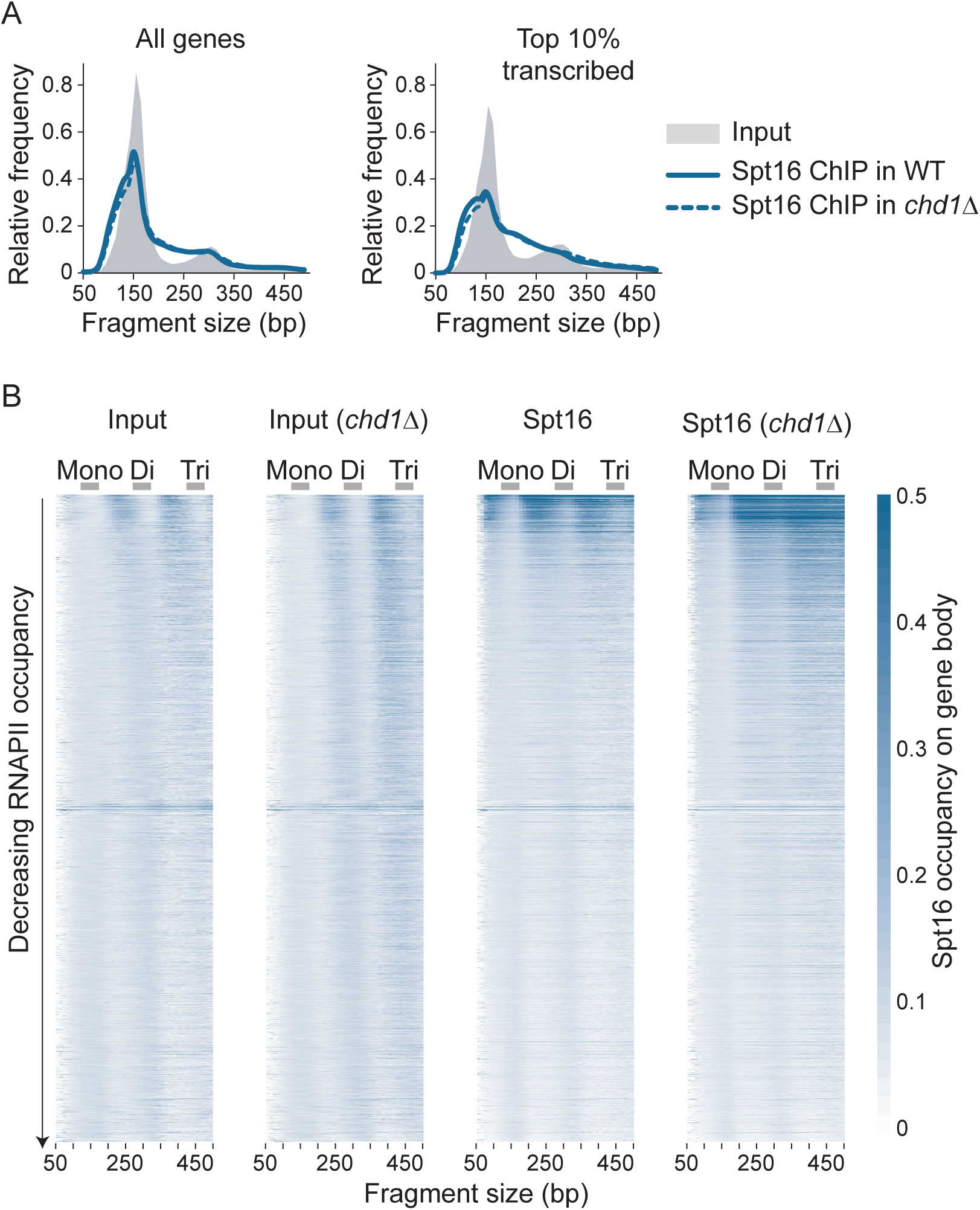
FACT (Spt16) binds disorganized nucleosomal particles on transcribed genes. **A**) Distribution plots of the size of DNA fragments recovered from Input (grey area) and FACT (Spt16, blue traces) MNase-ChIP-seq samples from WT (solid traces) and *chd1*Δ (dotted traces) cells for all genes (n = 5,796, left panel) and the top 10% transcribed genes (n = 580, right panel). **B**) Heatmap representation of Input and FACT (Spt16) average occupancy on gene body (n = 5,796, sorted by decreasing RNAPII occupancy on the y-axis) from WT and *chd1*Δ cells, as determined by MNase-ChIP-seq, computed with DNA fragments from different sizes (x-axis). Mono-, di- and tri-nucleosome-sized DNA fragments are indicated. See also **Figure S3**.

To systematically investigate the relationship between DNA fragment size associated with FACT and transcription, we generated heatmaps of the average fragment density over genes sorted by decreasing RNAPII occupancy (y-axis) for DNA fragments from different sizes (x-axis). **Figure 3B** clearly shows that the transcription-dependent signal in the Spt16 ChIP emanates from the internucleosomal size (150-300 bp and 300-450 bp) and subnucleosomal size (<150 bp) fragments, while the mononucleosomal or dinucleosomal size fragments, despite being more abundant in the sample (see distribution plots in **Figure 3A**), do not preferentially map to transcribed genes (similar results were obtained for Pob3, see **Figure S3B**). Hence, transcription-dependent FACT occupancy is reflected in the internucleosomal and subnucleosomal size fragments in these datasets. The mononucleosomal size fragments, which represent most of the signal (although depleted relative to the Input samples), may represent FACT binding to nucleosomes in other (non-transcriptional) contexts or, more likely, represent background signal in the ChIP. Taken together, these results show that FACT associates with disorganized nucleosomes present in transcribed regions.

### FACT spreads inside the gene body from the +1 nucleosome in a Chd1-dependent manner

To determine where FACT-associated DNA fragments map relative to genes, we generated two-dimensional occupancy (2DO) plots (Chereji et al., 2018; Henikoff et al., 2011) of the center of Input-and Spt16-associated fragments relative to the dyad of the +1 nucleosome in both WT and *chd1*Δ cells (**Figures 4A** and **4B**; see **Figures S4A** and **S4B** for similar experiments performed with Pob3). As expected, the Input samples are dominated by nucleosomal size fragments (150 bp) aligning with nucleosome dyads, with some dinucleosomal size fragments (300 bp) aligning between nucleosomal dyads (**Figure 4A**, top left panel). In addition to these dominant features, the Spt16 ChIPs from WT cells contain additional signals (notably between 150 bp and 300 bp) (**Figure 4A**, top right panel) which we showed to be the relevant signal for transcription-dependent FACT occupancy (see above). Interestingly, the plot revealed that the center of these internucleosomal size fragments covers the entire range of positions, forming an “inverted-v” linking the mononucleosomal and dinucleosomal signals. This result suggests that FACT-bound particles can occupy a wide range of positions relative to canonical nucleosomal locations and that the position of these fragments is linked to their size (see **Figure 4C** for a graphical representation of representative MNase-resistant DNA fragments observed in FACT ChIP samples). Fragments forming the upward diagonal of the inverted-v share a common 5’ extremity aligning with the upstream edge of their corresponding nucleosome but are increasingly longer from their 3’ end ascending the diagonal (**Figure 4C**, left panel, fragments *a*-*d*). These fragments therefore represent nucleosomal particles with extended MNase protection on the side distal to the transcription start site (TSS-distal). Conversely, the fragments along the downward diagonal have fixed 3’ ends aligning with the downstream edge of their nucleosomes and variable 5’ extremities (**Figure 4C**, right panel, fragments *d*-*g*). These fragments hence represent nucleosomal particles with extended MNase protection on the TSS-proximal side. In line with the analyses shown in **Figure 3**, these internucleosomal size fragments are more prominent on the most highly transcribed genes (**Figure 4A**, compare bottom-right panel with top-right panel). We surmise those fragments represent intermediate states of FACT − likely together with other factors− dynamically interacting with both sides of nucleosomes as they are being transcribed.

**Figure 4:**
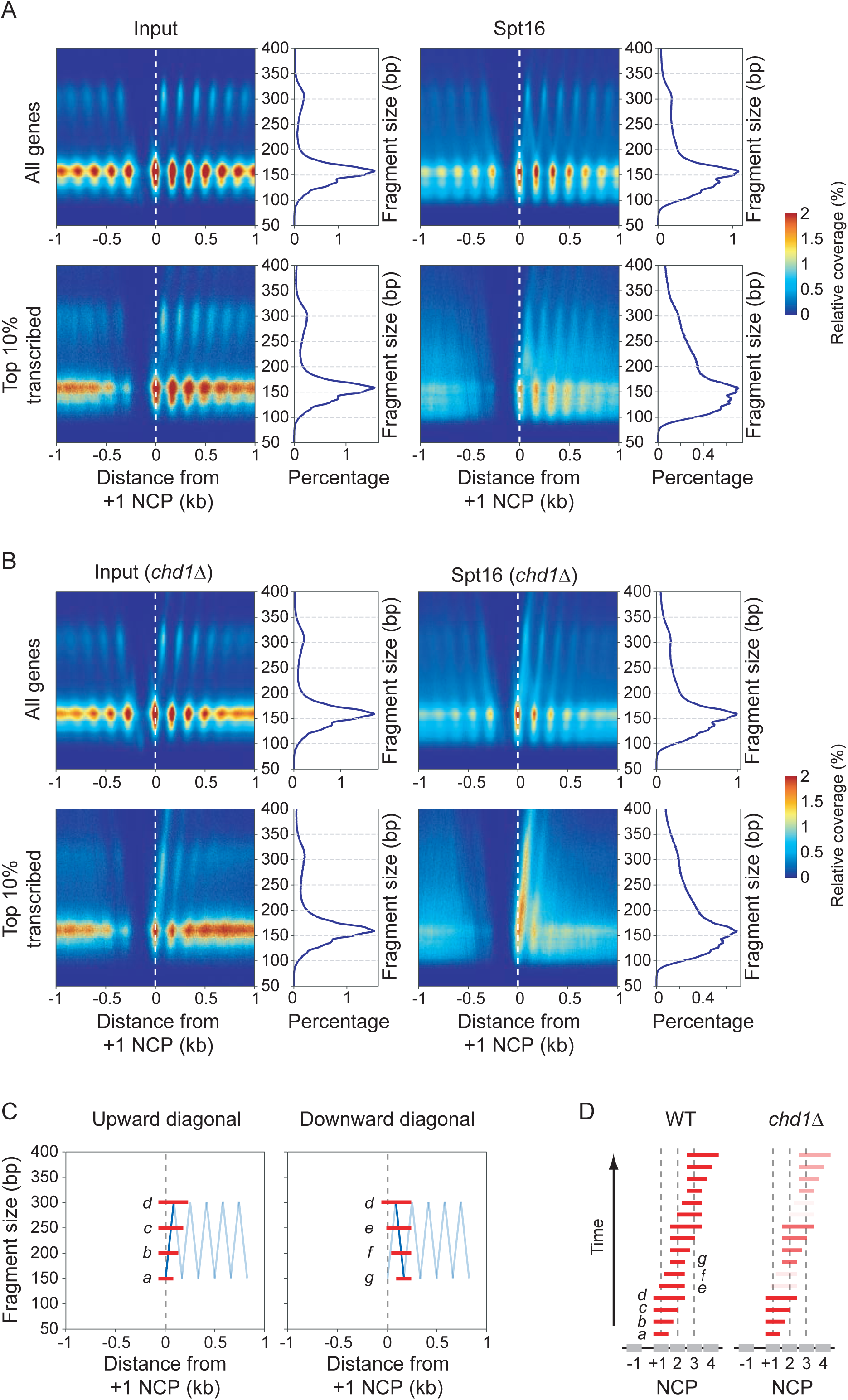
FACT spreads inside the gene body from the +1 nucleosome in a Chd1-dependent manner. **A**) 2DO plots of the coverage of the sequenced fragment mid-points (nucleosome dyads) in Input (left) and Spt16 ChIP (right) from MNase-digested chromatin from WT cells, relative to the +1 nucleosome dyads. On the right of each heatmap is the distribution of the fragment sizes. Top panels show data for all genes (n = 5,796) and bottom panels show data for the most transcribed genes (n = 580, top 10%). **B**) Same as panel A but for *chd1*Δ cells. **C**) A graphical representation of the fragments aligning of the inverted-v’s observed in 2DO plots from FACT ChIP samples. The blue diagonal lines highlight the upward and downward trajectories of the fragment mid-points for all fragments ranging from 150 bp to 300 bp starting from the +1 nucleosome to the +1/+2 dinucleosome (left) and from the +1/+2 dinucleosome to the +2 nucleosome (right) in the WT sample. Light blue lines depict the trajectory of the further downstream fragments. The red bars labeled “*a*” through “*d*” depict examples of fragments at different positions along the upward diagonal and the red bars labeled “*d*” through “*g*” depict examples of fragments at different positions along the downward diagonal. **D**) A working model for FACT spreading down a gene based on the MNase-ChIP-seq data in WT and *chd1*Δ cells from panel A. Fragments are considered as different snapshots of a dynamic process (y-axis speculatively denoted as “Time”) and were ordered as they appear when walking through the inverted-v zig-zag. Note the under-represented fragments and the progressive loss of signal in *chd1*Δ. See also **Figure S4**.

Interestingly, both Spt16 and Pob3 ChIPs from *chd1*Δ cells yielded dramatically different patterns (**Figure 4B** for Spt16 and **Figure S4B** for Pob3). First, and in agreement with our ChIP-chip and ChIP-exo data, the MNase-ChIP-seq signal decreases from 5’ to 3’ confirming that FACT does not distribute evenly along genes in the absence of the chromatin remodeler. Second, DNA fragments along the downward diagonal of the inverted-v (fragments *e* and *f* in **Figure 4C**) are absent, suggesting that in *chd1*Δ cells, FACT mostly binds nucleosomal particles with extended MNase protection on the TSS-distal side. Putting these observations together, we hypothesized that the different sized fragments represent intermediate states of a dynamic process by which FACT interacts with nucleosomes from the TSS-proximal to distal sides (**Figures 4C** and **4D**). The loss of the downward inverted-v fragments reflecting binding to the downstream side of the nucleosome in *chd1*Δ cells suggests Chd1 is required for the appearance of the later stage intermediates. Also, because FACT signal logarithmically decreases at each nucleosome from 5’ to 3’ in *chd1*Δ cells (**Figure S4C**), the data suggest that Chd1 allows FACT to translocate from one nucleosome to the next (see **Figure 4D**). Both in yeast (**Figure 2C**) and *Drosophila* (Skene et al., 2014), catalytically dead Chd1 is stranded at the 5’ end of genes. We, therefore, propose that Chd1 uses its remodeling activity to translocate down the gene and this enables FACT to spread downstream (**Figure 4D**).

To gain more insights about the mechanism, we performed MNase-ChIP-seq analyses of Chd1 (WT and catalytic dead mutant (K407R)). Similar to FACT, Chd1 ChIP from MNase enriches fragments of internucleosomal sizes that correlate with transcription (**Figures S3C** and **S3D**). Interestingly, 2DO plots of these fragments suggest that Chd1 binds (protects from MNase) the TSS-distal side of nucleosomes (**Figures S4D** and **S4E**). Consistent with our ChIP-chip data, the signal is concentrated on the +1 nucleosome in *chd1*-K407R cells (**Figure S4E**). These results are consistent with biochemical and structural data (Farnung et al., 2017; Sundaramoorthy et al., 2018) and suggest possible mechanisms for how Chd1 may contribute to FACT movement (See Discussion).

### A mathematical model captures Chd1-dependent spreading of FACT from the +1 nucleosome

Our model, although compatible with our and published data, assumes that the different MNase-protected fragments bound by FACT represent intermediates of a dynamic process. It also assumes that their appearance over time follows their appearance along the inverted-v patterns (see placement of fragments in **Figure 4D**). A formal validation of this model would require experimental evidence for the time-dependence of the appearance of the MNase-protected fragments. Such experiments, however, are not practically doable, notably because yeast genes are very short and no efficient methods would allow us to synchronize transcription with the sufficient time scale (yeast transcription can not be synchronized using RNAPII elongation inhibitor DRB (5,6-dichloro-1-β-D-ribofuranosylbenzimidazole) as commonly done in higher eukaryotes (Jonkers and Lis, 2015)). To challenge our model against alternative ones, we turned to mathematical modeling. We built stochastic models with multiple potential mechanisms (**Figures 5** and **S5**). Among the models tested, one (with a core “inchworm-like” mechanism of progression) was able to capture many qualitative features of the inverted-v pattern observed in the FACT MNase-ChIP data and can be summarized as follows. FACT binds at the +1 nucleosome location; extends from the downstream end in the direction of transcription; reaches full extension when encompassing the +1 and +2 nucleosome locations; transitions to a retracting state; retracts from the upstream end in the direction of transcription; reaches full retraction when encompassing the location of the +2 nucleosome; transitions to an extending state; resumes extension and repeats the extension/retraction process until the end of the gene is reached and the complex unbinds from chromatin (**Figure 5A**). Upon simulations, this model reached a steady-state recapitulating the characteristic inverted-v pattern observed in WT cells (**Figure S5A**). Adding a moderate rate of FACT unbinding at any point in the process was able to capture a shallow decrease in overall signal density towards the 3’ end of genes (**Figure S5B**). Implementing a relatively low transition rate from the extending to retracting states, and vice-versa, captured the additional density at the apices of the inverted-v pattern seen in the FACT MNase-ChIP data (**Figure S5C**). This refined model (**Figure 5B** and **Movie S1**) recapitulates the key features of the observed FACT MNase-ChIP in WT cells. We tested alternative models, but none were as successful in recapitulating the pattern observed in the FACT MNase-ChIP data (**Figures 5C, S5D-S5F**). Notably, allowing FACT to bind at any nucleosome location results in increased density towards the 3’ end of a gene, which is not observed in the MNase-ChIP data either from WT or *chd1*Δ cells, suggesting that binding of FACT predominantly happens at the beginning of a gene (**Figure 5C**).

**Figure 5:**
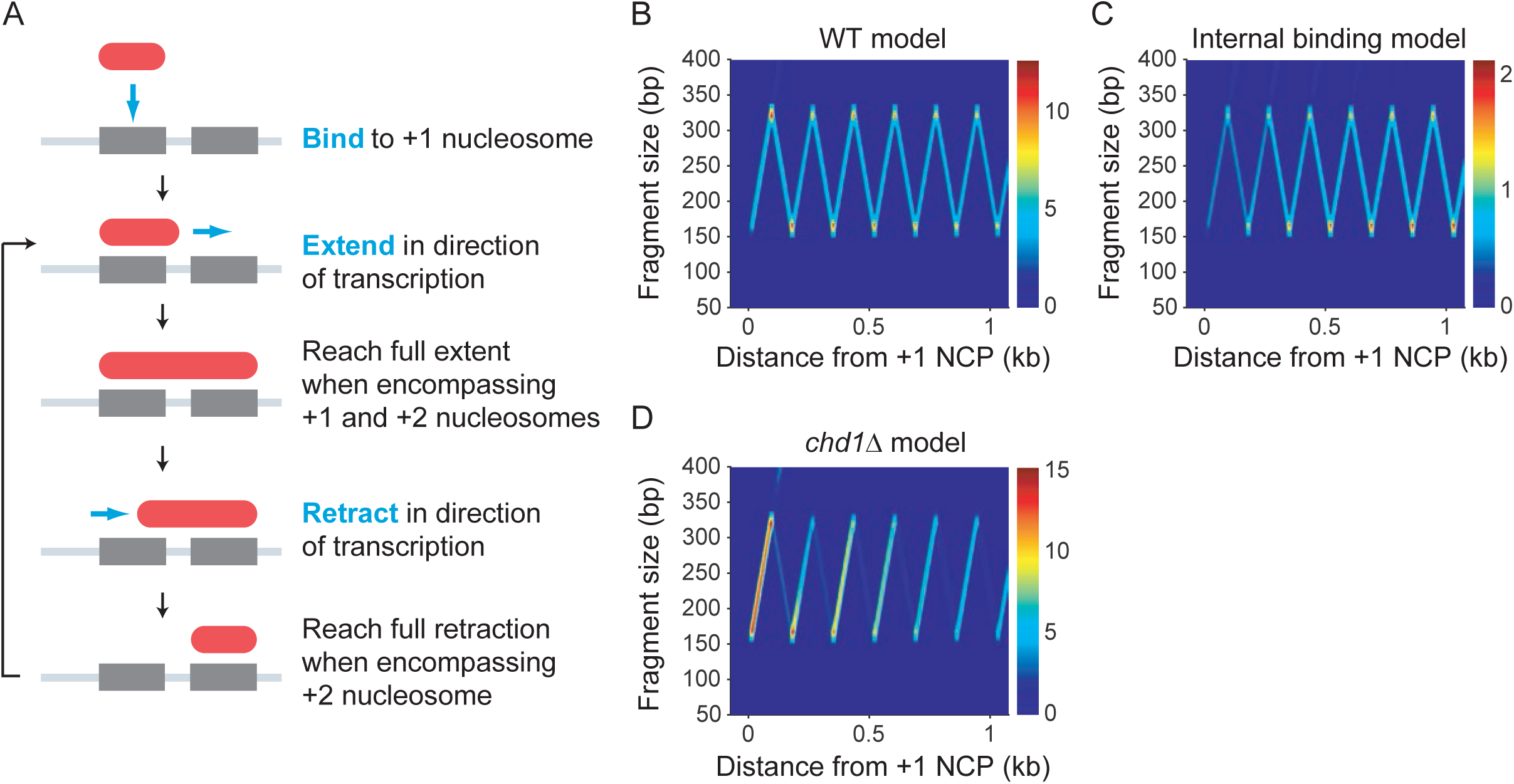
Mathematical modeling supports FACT binding at the +1 nucleosome and traversing the gene with an “inchworm” mechanism. **A**) Schematic of the inchworm mechanism. **B**) Simulated protected fragment sizes and locations from the basic model with constant unbinding and reduced transition rates. **C**) Simulated protected fragment sizes and locations from an alternative model with binding of FACT allowed at any nucleosome location within a gene. **D**) Simulated protected fragment sizes and locations for the same model as in A) but with a reduced rate of extension. The units of the color bars in panels B-D represent the number of fragments observed in a complete simulation averaged over a 20 ⨯ 20 bp window (⨯ 10^5^). See also **Figure S5**.

We then took advantage of our mathematical model to test possible mechanisms for the role of Chd1. Our ChIP-chip data show that Chd1 contributes to the distribution of FACT via its ATPase activity (**Figure 2C**). Hence, we envisioned that Chd1 may impact on any step in the FACT dynamic process. We therefore systematically tested the impact of increasing or decreasing each variable in the model (extension, transition, retraction, unbinding; see **Figures 5D, S5D-S5F**). Among those simulations, impairing the extension rate recapitulated all features of the *chd1*Δ experimental data: i) enhanced signal on the upward diagonal of the inverted-v pattern, ii) loss of the downward diagonal of the inverted-v and iii) a significant decrease in intensity towards the 3’ end of the gene (**Figure 5D** and **Movie S2**). Since in our mathematical model impairing the extension rate was the only change from the WT situation that recapitulates the *chd1*Δ pattern, we can confidently propose that Chd1 promotes FACT spreading by energetically favoring the extension step, which −in biochemical terms− represents extending the MNase protection downstream of the nucleosome.

In the course of building and testing our models, we noticed that impairing the extension rate (a condition mimicking the *chd1*Δ data) systematically led to the appearance of increased signal for fragments greater than 300 bp as a natural consequence of not allowing FACT complexes to overlap (**Figure S5G**, left panel). Looking back at the experimental data, we noticed that such “higher-order” signal was also observed in the *chd1*Δ MNase-ChIP data when displaying fragments up to 800 bp (**Figure S5G**, right panel). Because these fragments were not used in the construction of the mathematical model, their presence in both the simulations and the experimental data represents an internal validation of the model. In summary, our mathematical modeling data fully support a mechanism where FACT enters a gene by binding to the +1 nucleosome and spreads downstream using a cycle of steps that translate into successive extensions and retractions of the MNase-protection of the nucleosomes, with the help of Chd1. The model also suggests that Chd1 contributes most to the first (extension) step of this process.

### FACT recognizes partially unwrapped +1 nucleosomes induced by RNAPII

While the data presented above provide a mechanism to explain how FACT spreads down a gene after being recruited to the +1 nucleosome, it does not explain how it is initially recruited to the +1 nucleosome of transcriptionally active genes. We addressed this question by analyzing the subnucleosomal-size DNA fragments from FACT MNase-ChIP-seq experiments. We mapped the center of DNA fragments of different sizes on the dyad of the +1 nucleosome. As previously shown in Drosophila (Ramachandran et al., 2017), the distribution of subnucleosomal-size DNA fragments from MNase-seq is bimodal and can be fit to two Gaussians, representing two populations of nucleosomes partially unwrapped from one end or the other (see **Figure 6A** for a graphical representation). Hence, guided by this previous work (Ramachandran et al., 2017), we mapped the distributions of fragments of 90 bp, 103 bp, and 125 bp around the dyad of the +1 nucleosome both in WT and *chd1*Δ cells. The 90 bp and 103 bp fragments informing on asymmetrically unwrapped nucleosomal particles and the 125 bp fragments reflecting symmetrically unwrapped particles (Ramachandran et al., 2017). Looking at the Input material of WT cells revealed that the proportion of +1 nucleosomes partially unwrapped from the TSS-proximal side is similar to those unwrapped from the TSS-distal side, suggesting asymmetric nucleosome breathing (**Figures 6B** and **S6A**, 103 bp and 90 bp fragments plots for Input). As previously described (Ramachandran et al., 2017), symmetrical unwrapping is also observed, as shown through the analysis of the 125 bp fragments (**Figure 6B**). Also note that similar observations were made when looking at the +2 nucleosome (**Figure S6B**). Subnucleosomal-size DNA fragments from the FACT ChIPs, however, were heavily biased for asymmetrically unwrapping from the TSS-proximal side (**Figures 6B** and **S6A**, 103 bp and 90 bp fragments plots for Spt16 and Pob3 ChIP) demonstrating that FACT preferentially binds to +1 nucleosomes with TSS-proximal DNA contact loss. Interestingly, similar results were observed in *chd1*Δ cells, further supporting the idea that the chromatin remodeler is involved in post-recruitment steps (FACT spreading) rather than during the initial recruitment at the +1 nucleosome. Also noteworthy is that the enrichment of particles with TSS-proximal unwrapping was not observed to the same extent at the +2 nucleosome (**Figure S6B**), again suggesting that FACT engages the +1 nucleosome differently than it does with downstream ones. Hence, we propose that transcription-induced unwrapping of the TSS-proximal side of the +1 nucleosome triggers FACT recruitment.

**Figure 6:**
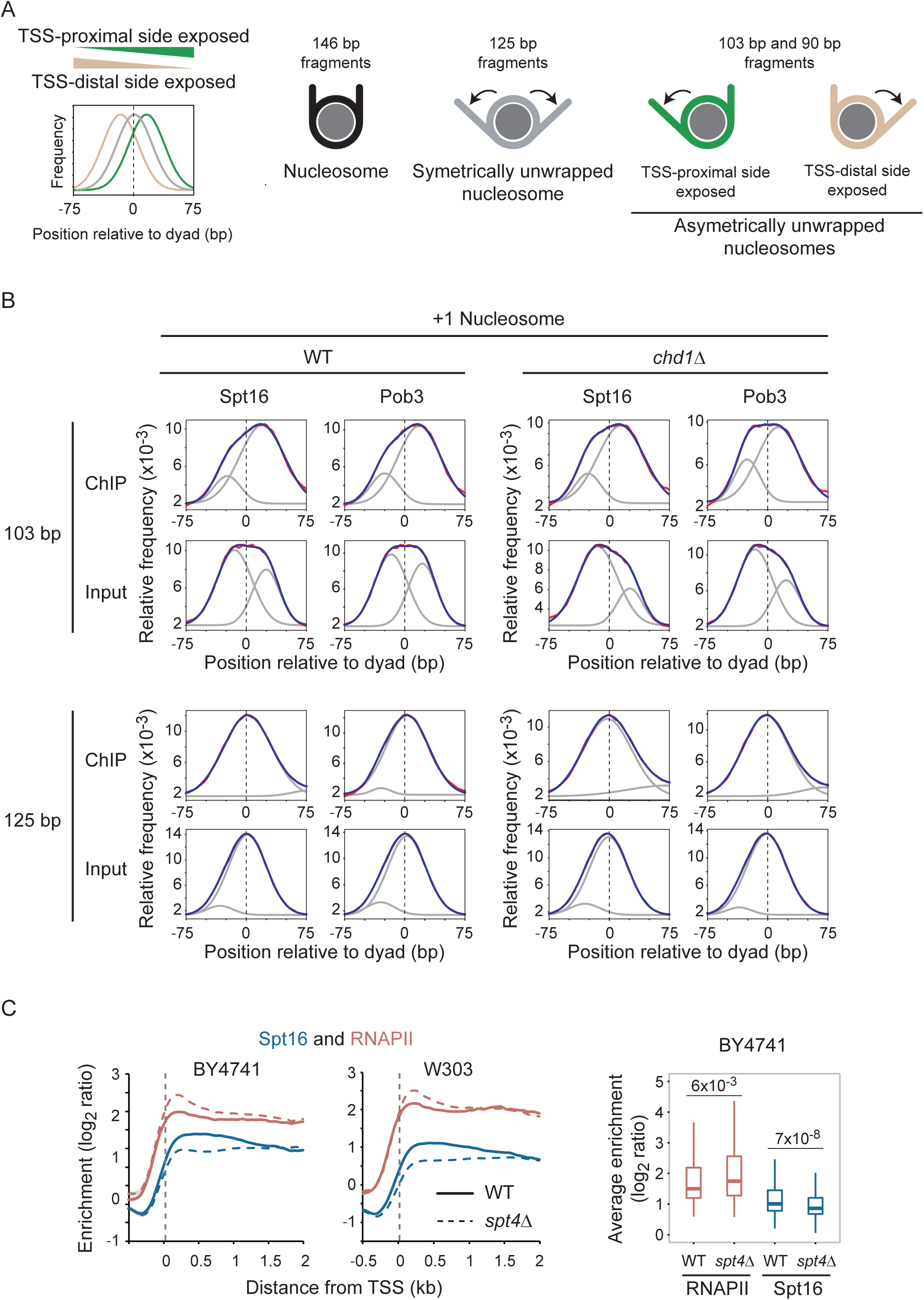
FACT recognizes +1 nucleosomes, asymmetrically unwrapped from their TSS-proximal side. **A**) Graphical explanation for the interpretation of the distribution plots shown in panel B. The graphs show the distribution of the center of subnucleosomal size fragments. A distribution centered at “0” (grey trace) indicates symmetrically unwrapped nucleosomal particles, whereas a distribution centered to the left (tan trace) or the right (green trace) indicates nucleosomal particles asymmetrically unwrapped on their TSS-distal and TSS-proximal side, respectively (see cartoons on the right with respective colors). We used 90 bp, 103 bp, and 125 bp fragments as per (Ramachandran et al., 2017). **B**) Distribution of 103 and 125 bp (+/-5 bp) fragment centers (red), from Spt16 and Pob3 MNase-ChIP-seq experiments (and their Inputs) from WT and *chd1*Δ cells, plotted relative to the position of +1 nucleosome dyads. The data (red) were fitted to a double-Gaussian (blue). The grey traces show the two individual Gaussians. **C**) Full recruitment of FACT requires Spt4/Spt5. Left: Metagene of FACT (Spt16) and RNAPII (Rpb3) occupancy over transcribed genes (n = 275), as determined by ChIP-chip, relative to Input, in WT (solid traces) and *spt4*Δ (dashed traces) cells. Data are shown for experiments performed in two genetic backgrounds (BY4741 and W303). Right: A box plot of the average RNAPII and Spt16 occupancy in WT and *spt4*Δ cells for transcribed genes (n = 275) from the BY4741 experiment. See also **Figure S6**.

The analysis presented above supports the idea that RNAPII mediated unwrapping of the +1 nucleosome triggers the initial recruitment of FACT in the 5’end of transcribed genes. This model is consistent with several *in vitro* observations showing that while FACT does not bind intact nucleosomes, it binds to nucleosomes that are partially unwrapped by several means (see Introduction). Further support for that model came from our FACT ChIP-chip experiments in *spt4*Δ cells. Spt4/Spt5 (also known as DSIF in higher eukaryotes) is an elongation factor that interacts with FACT (Foltman et al., 2013; Han et al., 2010; Lindstrom et al., 2003). Detailed *in vitro* analyses showed that Spt4/5 stabilizes a partially unwrapped state of the nucleosome during transcription *in vitro* (Crickard et al., 2017). In another study, cryo-EM revealed that Spt4/5 prevents DNA from reassociation with histones when RNAPII is stalled near the dyad (Ehara et al., 2019). Together, these two studies support a role for Spt4/5 in assisting RNAPII negotiating with nucleosomes by preventing undesirable DNA-histone interactions that may cripple the progression of the polymerase through nucleosomes, notably nucleosome +1. Consistent with these *in vitro* experiments, our ChIP-chip analyses of RNAPII in *spt4*Δ cells revealed an accumulation at the 5’ end of genes (**Figure 6C**). Interestingly, FACT occupancy is decreased in *spt4*Δ cells (**Figure 6C**). This observation contrasts with other mutants we have tested (see above), where variations in RNAPII occupancy are generally mirrored by FACT. The decreased FACT recruitment in a context where RNAPII is accumulated rather suggests that Spt4/5 contributes to FACT recruitment. In light of the *in vitro* data mentioned above (Crickard et al., 2017), we propose that Spt4/5 contributes to the unwrapping (by preventing unsuitable rewrapping) of the +1 nucleosome, hence contributing to the proper recruitment of FACT. In the absence of Spt4/5, the unwrapping/rewrapping process is crippled, and undesirable DNA-histone contacts are established both blocking polymerase progression and preventing FACT recruitment.

## DISCUSSION

How FACT is recruited to genes is a long-standing question that remained elusive. Here, we provide an in-depth analysis of FACT recruitment in *S*. *cerevisiae*. Using combinations of ChIP-chip, ChIP-exo, and MNase-ChIP-seq in WT and various mutants, we show that FACT binds transcribed chromatin, rather than RNAPII, in agreement with a recent report (Martin et al., 2018). Furthermore, our work supports FACT binding asymmetrically unwrapped nucleosomes at the 5’ end of genes as they are being approached by the polymerase. We also show that the proper distribution of FACT along genes requires the ATPase activity of the chromatin remodeler Chd1. Mathematical modeling reveals that Chd1 stimulates the initial steps of a stepwise, inchworm-like process allowing FACT to move from one nucleosome to the next. Hence, our data suggest that FACT works via a processive mechanism on the chromatin template as it is transcribed and remodeled by RNAPII and Chd1.

FACT recognizing partially unwrapped nucleosomes is consistent with a plethora of *in vitro* FACT-nucleosome interaction data (Formosa et al., 2001; Liu et al., 2020; McCullough et al., 2018; Nesher et al., 2018; Ruone et al., 2003; Tsunaka et al., 2016; Wang et al., 2018). *In vivo*, partially unwrapped nucleosomes were detected in transcribed regions in Drosophila (Ramachandran et al., 2017) and yeast (this study). In addition, recent structural and biochemical studies showed DNA unwrapping from the upstream side of RNAPII-engaged nucleosomes (Farnung et al., 2018; Kujirai et al., 2018). It is therefore reasonable to think that asymmetrical DNA unwrapping of the +1 nucleosome −triggered by RNAPII− would promote FACT recruitment. Because such partial unwrapping can be achieved multiple ways and in different circumstances in cells, the model proposed here −although elaborated looking at FACT in the context of RNAPII transcription− provides an attractive recruitment mechanism for FACT to function in different contexts. Indeed, although FACT was originally identified as a factor assisting RNAPII elongation through nucleosomes (Belotserkovskaya et al., 2003; Orphanides et al., 1998), it is involved in many nuclear processes including transcription by all three nuclear RNA polymerases, DNA replication and DNA repair (reviewed in (Formosa, 2013; Gurova et al., 2018)). In addition, FACT also works at some promoters, where it helps in transcription factor and coactivator binding and nucleosome redeposition (Biswas et al., 2006; Biswas et al., 2005; Takahata et al., 2009; Voth et al., 2014). Because nucleosomes need to unwind during all these processes, we propose that nucleosome unwrapping (mediated by different activities in each context) is triggering FACT recruitment in these settings as well. Alternative models, such as direct recruitment of FACT by the different machineries, appear unlikely as it would require FACT to specifically interact with several proteins; yet FACT is essentially composed of domains dedicated to histones and DNA interactions (reviewed in (Zhou et al., 2020)).

In addition to its role during elongation, FACT had been shown to play roles in initiation at some inducible genes (Biswas et al., 2006; Biswas et al., 2005; Takahata et al., 2009; Voth et al., 2014). Also, recent studies showed that RNAPII (Jeronimo et al., 2019; Petrenko et al., 2019) and TFIIB (Petrenko et al., 2019) occupancies are dramatically decreased at active genes when FACT subunits are disrupted. While these results may suggest a direct and global role for FACT in initiation, our data suggest otherwise. Indeed, we ruled out recruitment of FACT at promoters genome-wide. Hence, although we cannot exclude (and do not dispute) roles at specific promoters, our data suggest that FACT impacts initiation genome-wide indirectly via its function during elongation where it prevents cryptic transcription and the consequent titration of the transcription machinery, as previously suggested for Spt6 (Doris et al., 2018). This scenario appears plausible in the light of the recent finding that RNAPII is limiting in yeast cells (Sun et al., 2020)

Our work shows that proper distribution of FACT along genes requires the catalytic activity of Chd1. Because tampering with Chd1 affects FACT distribution but not its occupancy levels, we propose that, once recruited on chromatin at the 5’ end of genes, FACT translocates towards the 3’ end of genes with the help of Chd1. Based on MNase-ChIP-seq data and mathematical modeling, we propose that this translocation involves successions of gain and loss of contacts between FACT, histones, and DNA. The translocation of FACT is therefore likely to be coupled to its chaperone activity; that is, as it “chaperones” histones during RNAPII progression through a nucleosome, FACT makes its way from one nucleosome to the next. Our data suggest that during this process, FACT initially makes contact with a single nucleosome and progresses towards intermediates where it ultimately contacts two successive nucleosomes before letting go of the initial one. Interestingly, our mathematical modeling revealed an important role for Chd1 in the first part of this “inchworm-like” process, that is, transiting from binding to one nucleosome into simultaneously contacting two nucleosomes. It is tempting to speculate that this capacity to bind successive nucleosomes may contribute to FACT’s ability to maintain proper histone modification patterns during transcription and (perhaps) replication (Jamai et al., 2009; Jeronimo et al., 2019; Jeronimo et al., 2015).

How does Chd1 remodeling activity promote FACT translocation? Consistent with our MNase-ChIP-seq data, structural studies showed that Chd1 makes contacts with downstream DNA (Farnung et al., 2017; Sundaramoorthy et al., 2018). This unique feature among remodelers is achieved through its SANT and SLIDE domains. In addition, the ATPase domain contacts DNA two superhelical turns upstream of the dyad on the TSS-proximal side and “pumps” DNA towards the dyad. This creates forces that should favor the breakage of histone-DNA contacts around the dyad and propagate DNA towards TSS-distal side. It is tempting to speculate that this process is driving FACT progression forward.

We surmise that the recently described structure of FACT binding a partially unwrapped nucleosome (Liu et al., 2020) represents the initial state of FACT binding chromatin. It is worth pointing out that the “di-nucleosome footprints” we observe in our FACT MNase-ChIP-seq data are likely to include other proteins besides FACT and histones (including Chd1, RNAPII, and other elongation factors). It is hence impossible, from our *in vivo* data alone, to determine whether FACT directly contacts successive nucleosomes. Our findings therefore call for the investigation of structures of FACT bound to more complex substrates (e.g. di-nucleosomes) together with Chd1, RNAPII, and perhaps other elongation factors. Several cryo-EM structures have recently been solved that should help in that venture (Ehara et al., 2019; Ehara et al., 2017; Farnung et al., 2018; Farnung et al., 2017; Kujirai et al., 2018; Liu et al., 2020; Vos et al., 2018; Vos et al., 2020; Xu et al., 2017). Collectively, the work presented here ties previous *in vitro* information to a relevant *in vivo* context (transcription) and provides a ground state for the dissection of FACT chaperoning mechanisms, both *in vivo* and *in vitro*.

## ACKNOWLEDGMENTS

We are grateful to N. Francis for critical reading of the manuscript. We also thank A. Bataille for advice about ChIP-exo library preparation and analysis; M. Sarsenova for generating strains yFR3061, yFR3084, yFR3075; O. Rocheleau-Leclair for generating strains yFR2024, yFR2035; and T. Formosa, R.A. Young, S. Hahn, S. Buratowski and F. Winston for sharing reagents. This work was funded by a grant from the Canadian Institutes of Health Research (CIHR) to F.R. (MOP-162334). This research was also enabled in part by support provided by « Calcul Québec » (http://www.calculquebec.ca) and Compute Canada (www.computecanada.ca). F.R. holds a Research Chair from the « Fonds de Recherche Québec –Santé » (FRQS). P.C. was supported by a studentship from the FRQS. A.A. was supported by a BBSRC grant to J.M. (BB/S009035/1).

## AUTHOR CONTRIBUTIONS

F.R. and C.J. conceived the study and designed experiments; C.J. performed most experiments; A.A. performed the mathematical modeling; P.C. performed some ChIP-chip experiments in Figures 1E and S2; C.P. and F.R. performed computational data analyses; F.R. supervised research. J.M. supervised modeling. F.R. and J.M provided funding. F.R. and C.J. wrote the manuscript with input from all authors.

## DECLARATION OF INTERESTS

The authors declare no competing interests.

## FIGURE LEGENDS

**Figure S1**: FACT occupancy correlates with transcription by RNAPII but that does not require CTD phosphorylation, related to **Figure 1. A**) Metagenes of RNAPII (Rpb3) and FACT (Spt16, Pob3) occupancy over highly (dark shade; n = 85), mildly (medium shade; n = 190) and lowly (light shade; n = 2,141) transcribed genes, as determined by ChIP-chip, relative to Input, in WT cells. **B**) Scatter plots of Spt16 versus RNAPII (Rpb3; left), Pob3 versus RNAPII (Rpb3; middle), and Pob3 versus Spt16 (right) occupancy on ORFs (n = 6,692), as determined by ChIP-chip, relative to Input, in WT cells. **C**) Metagenes of RNAPII (Rpb3) and FACT (Spt16, Pob3) occupancy over transcribed genes (n = 275), as determined by ChIP-chip, relative to Input. Top panels, WT (solid traces), *cdk8*Δ (dashed traces) and *bur2*Δ (dotted traces) cells; Middle panels, WT (solid traces) and *ctk1*Δ (dashed traces) cells; Bottom panels, WT (solid traces), *kin28*as (dashed traces) and *kin28*as/*bur2*Δ (dotted traces) cells. WT, *kin28*as, and *kin28*as/*bur2*Δ cells in the bottom panels were treated 15 min with NAPP1.

**Figure S2**: FACT recruitment is largely independent of PAF1C, the capping enzyme, chromatin modifiers, and chromatin remodelers but affected in *chd1*Δ cells, related to **Figure 2. A**) Metagenes of FACT (Spt16) and RNAPII (Rpb3) occupancy over transcribed genes (n = 275), as determined by ChIP-chip, relative to Input, in WT (solid traces) and PAF1C mutant (*paf1*Δ, *leo1*Δ, *ctr9*Δ, *cdc73*Δ, *rtf1*Δ; dashed traces) cells. **B**) Metagenes of FACT (Spt16) and RNAPII (Rpb3) occupancy over transcribed genes (n = 275), as determined by ChIP-chip, relative to Input, in WT (solid traces) and *cet1*ΔN (dashed traces) cells. **C**) Metagenes of FACT (Spt16) and RNAPII (Rpb3) occupancy over transcribed genes (n = 275), as determined by ChIP-chip, in WT (solid traces), sas3Δ, *gcn5*Δ, *set2*Δ, *bre1*Δ, *ubp8*Δ and *ubp10*Δ (dashed traces) cells. **D**) Metagenes of FACT (Spt16) and RNAPII (Rpb3) occupancy over transcribed genes (n = 275), as determined by ChIP-chip, relative to Input, in WT (solid traces) and *chd1*Δ (dashed traces) cells from W303 and BY4741 genetic backgrounds. **E**) Metagenes of FACT (Spt16) and RNAPII (Rpb3) occupancy over transcribed genes (n = 275), as determined by ChIP-chip, relative to Input, in WT (solid traces), *isw1*Δ (dashed traces, top panel) and *isw2*Δ (dashed traces, bottom panel) cells. **F**) Metagenes of histones H2B and H4, RNAPII (Rpb3) and FACT (Spt16, Pob3) occupancy over transcribed genes (n = 275), as determined by ChIP-chip, relative to Input, in WT (solid traces) and *spt6-1004* (dashed traces) cells which were shifted to 37°C for 80 min. Data for H2B and H4 are from (Jeronimo et al., 2019) and (Jeronimo et al., 2015), respectively.

**Figure S3**: Analysis of the size of the mapped DNA fragments from the FACT (Pob3) and Chd1 MNase-ChIP-seq experiments, related to **Figure 3. A**) Distribution plots of the size of DNA fragments recovered from Input (grey area) and Pob3 (violet traces) MNase-ChIP-seq samples from WT (solid traces) and *chd1*Δ (dotted traces) cells. Top panel shows data for all genes (n = 5,796) and bottom panel shows data for the top 10% transcribed genes (n = 580) genes. **B**) Heatmap representation of Input and Pob3 average occupancy on gene body (n = 5,796, sorted by decreasing RNAPII occupancy on the y-axis) from WT and *chd1*Δ cells, as determined by MNase-ChIP-seq, computed with DNA fragments from different sizes (x-axis). **C**) Distribution plots of the size of DNA fragments recovered from Input (grey area) and Chd1 (black traces) MNase-ChIP-seq samples from WT (solid traces) and *chd1*-K407R (dotted traces) cells. Top panel shows data for all genes (n = 5,796) and bottom panel shows data for the top 10% transcribed genes (n = 580) genes. **D**) Heatmap representation of Input and Chd1 average occupancy on gene body (n = 5,796, sorted by decreasing RNAPII occupancy on the y-axis) from WT and *chd1*-K407R cells, as determined by MNase-ChIP-seq, computed with DNA fragments from different sizes (x-axis). Mono-, di-and tri-nucleosome-sized DNA fragments are indicated.

**Figure S4**: 2DO plots for Pob3 and Chd1 MNase-ChIP-seq experiments, related to **Figure 4. A**) 2DO plots of the coverage of the sequenced fragment mid-points (nucleosome dyads) in Input (left) and Pob3 ChIP (right) from MNase-digested chromatin from WT cells, relative to the +1 nucleosome dyads. On the right of each heatmap is the distribution of the fragment sizes. Top panels show data for all genes (n = 5,796) and bottom panels show data for the most transcribed genes (n = 580, top 10%). **B**) 2DO plots of the coverage of the sequenced fragment mid-points (nucleosome dyads) in Input (left) and Pob3 ChIP (right) from MNase-digested chromatin from *chd1*Δ cells, relative to the +1 nucleosome dyads. On the right of each heatmap is the distribution of the fragment sizes. Top panels show data for all genes (n = 5,796) and bottom panels show data for the most transcribed genes (n = 580, top 10%). **C**) Violin plot of the median signal from Spt16 MNase-ChIP-seq in *chd1*Δ cell (data shown in **Figure 4B**) over the six first nucleosomes for the top 10% transcribed genes (n = 580). The occupancies are calculated for all fragments in the 200-300 bp interval and using a nucleosome repeat length of 161 bp to delineate nucleosome territories. Red bars indicate the median. **D**) 2DO plots of the coverage of the sequenced fragment mid-points (nucleosome dyads) in Input (left) and Chd1 ChIP (right) from MNase-digested chromatin from WT cells, relative to the +1 nucleosome dyads. On the right of each heatmap is the distribution of the fragment sizes. Top panels show data for all genes (n = 5,796) and bottom panels show data for the most transcribed genes (n = 580, top 10%). **E**) 2DO plots of the coverage of the sequenced fragment mid-points (nucleosome dyads) in Input (left) and Chd1 ChIP (right) from MNase-digested chromatin from *chd1*-K407R cells, relative to the +1 nucleosome dyads. On the right of each heatmap is the distribution of the fragment sizes. Top panels show data for all genes (n = 5,796) and bottom panels show data for the most transcribed genes (n = 580, top 10%).

**Figure S5**: Mathematical modeling indicated that many plausible mechanisms are not a good fit to the observed experimental data, related to **Figure 5. A**) The most basic version of the model with FACT binding at the +1 nucleosome location, extension, and retraction. **B**) The basic version of the model, as in A), with constant FACT unbinding. **C**) The basic version of the model, as in A), with reduced rates of transition between the extending and retracting states. **D**) The full version of the model as in **Figure 5A** but with mid-gene loading and constant unbinding. **E**) The version of the model as in D) but with higher unbinding. **F**) The full version of the model as in **Figure 5A** but with mid-gene loading, constant unbinding, and a reduced retraction rate. **G**) Left: The full version of the model with reduced extension to match the *chd1*Δ mutant (as in **Figure 5D**) but depicting with larger fragment sizes up to 800 bp. Right: The experimental data for *chd1*Δ cells (same as in **Figure 4B**, top right panel) but depicting with larger fragment sizes up to 800 bp. The units of the color bars in the simulation panels represent the number of fragments observed in a complete simulation averaged over a 20⨯20 bp window (⨯10^5^).

**Figure S6:** Distribution of subnucleosomal-size DNA fragments from FACT MNase-ChIP-seq experiments, relative to +1 and +2 nucleosome dyads, related to **Figure 6. A**) Distribution of 90 bp (+/-5 bp) fragment centers (red), from Spt16 and Pob3 MNase-ChIP-seq experiments (and their Inputs) from WT and *chd1*Δ cells, plotted relative to the position of +1 nucleosome dyads. The data (red) were fitted to a double-Gaussian (blue). The grey traces show the two individual Gaussians. **B**) Distribution of 90, 103, and 125 bp (+/-5 bp) fragment centers (red), from Spt16 and Pob3 MNase-ChIP-seq experiments (and their Inputs) from WT and *chd1*Δ cells, plotted relative to the position of +2 nucleosome dyads. The data was fitted to a double-Gaussian (blue). The grey traces show the two individual Gaussians.

**Movie S1**: Animation of the “inchworm-like” model of FACT recruitment spreading for the WT case. The left panel shows a representation of the regions of DNA protected from MNase digestion by FACT complexes for 100 realisations of a simulation across a synthetic gene. White bars represent FACT complexes, the x-axis is location in the gene and each track on the y-axis represents an individual realisation (tracks are separated by blank spaces). The right panel shows the density of fragments, of given sizes (y-axis) and locations (x-axis), protected from MNase digestion by FACT complexes as they are sampled from the simulation. Parameters of the simulation are as for the WT case as in the main text (**Figure 5B** and **Table 1**) except that the number of time steps shown is 10^5^, there are 100 realisations and there is no relaxation period prior to measurements for the protected fragments being taken.

**Movie S2**: Animation of the “inchworm-like” model of FACT recruitment spreading for the *chd1*Δ case (same as WT except for a reduced extension rate of the FACT complex). The left panel shows a representation of the regions of DNA protected from MNase digestion by FACT complexes for 100 realisations of a simulation across a synthetic gene. White bars represent FACT complexes, the x-axis is location in the gene and each track on the y-axis represents an individual realisation (tracks are separated by blank spaces). The right panel shows the density of fragments, of given sizes (y-axis) and locations (x-axis), protected from MNase digestion by FACT complexes as they are sampled from the simulation. Parameters of the simulation are as for the *chd1*Δ case as in the main text (**Figure 5D** and **Table 1**) except that the number of time steps shown is 10^5^, there are 100 realisations and there is no relaxation period prior to measurements for the protected fragments being taken.

## STAR METHODS

Detailed methods are provided in the online version of this paper and include the following: Outline to be created by the Production team.

## RESOURCE AVAILABILITY

### Lead Contact

Further information and requests for resources and reagents should be directed to and will be fulfilled by the Lead Contact, François Robert.

### Materials Availability

Yeast strains and plasmids generated in this study are available upon request.

### Data and Code Availability

- The microarray (ChIP-chip) and DNA sequencing (ChIP-exo and MNase-ChIP-seq) data generated during this study are available at the NCBI Gene Expression Omnibus (GEO; http://www.ncbi.nlm.nih.gov/geo/) under accession number GSE155144.
- The in-house scripts used to generate some of the analyses during this study are available at https://github.com/francoisrobertlab.
- The code related to the mathematical modeling of the MNase-ChIP-seq data is available at https://github.com/aangel-code/FACT-inchworm-model.

## EXPERIMENTAL MODEL AND SUBJECT DETAILS

*S*. *cerevisiae* strains used in this study are described in **Table S1**. Unless otherwise indicated, cells were grown at 30°C and 200 rpm in YPD (yeast extract-peptone-2% glucose, supplemented with 44 µM adenine) medium as follows. Strains were streaked from glycerol stocks onto 2% agar YPD plates and grown at 30°C for 2-3 days. An isolated colony was then grown overnight in 10 mL of YPD. This preculture was used to inoculate 50 mL of YPD at an OD_600_ of 0.1 which was grown to an OD_600_ of 0.7-0.9.

## METHOD DETAILS

### Yeast strains and plasmids

Genotypes for the yeast strains used in this study are listed in **Table S1**. HA-tag was inserted by homologous recombination of a PCR cassette amplified from the pMPY-3xHA plasmid (Schneider et al., 1995). The *chd1*-K407R catalytic dead point mutation (Simic et al., 2003) was introduced directly in the genome by CRISPR-Cas9 editing as previously described (Collin et al., 2019) using a pair of oligonucleotides corresponding to a gRNA targeting a region in Chd1 around amino acid K407 (see **Table S2**) designed for cloning into the pML107 plasmid (Laughery et al., 2015) and a PCR product obtained with repair oligonucleotides (see **Table S2**) containing the desired mutations (including one silent mutation destroying the protospacer adjacent motif and one creating the K407R mutation). Gene deletions were made by replacing the gene of interest with an auxotrophic or antibiotic resistance marker.

### Growth conditions

For shutting off transcription experiments, a strain harboring a temperature-sensitive allele of RNAPII (*rpb1-1*) was grown to an OD_600_ of 0.7 and an equal volume of YPD pre-warmed to 49°C was added to the flask, which was immediately transferred to 37°C for additional 80 min. For inhibition of Kin28 activity experiments, ATP analog-sensitive *kin28as* strains were precultured in yeast nitrogen base (YNB) medium lacking uracil before inoculation in YPD medium. *kin28as* strains and their controls were then grown to an OD_600_ of 0.8 and treated with 6 μM of 1-Naphthyl PP1 (NAPP1; Tocris Bioscience) for 15 min. Strains expressing the RPB1 CTD WT or CTD mutant plasmids were grown in YNB medium lacking histidine to an OD_600_ of 0.5 and treated with 1 μg/mL of rapamycin (Bio Basic) for 90 min to anchor-away the endogenous Rpb1 protein. For the experiments involving the *spt6-1004* temperature-sensitive allele, cells were grown to an OD_600_ of 0.5 and transferred to 37°C for 80 min.

### ChIP

ChIP experiments were performed from two independent biological replicates according to (Jeronimo et al., 2015). In brief, yeast cultures were grown in 50 mL of the appropriate medium (see above) to an OD_600_ of 0.7-0.9 before crosslinking with 1% formaldehyde (Fisher Scientific, BP531-500) at room temperature for 30 min and quenched with 125 mM glycine. Crosslinked cells were collected by centrifugation and washed twice with 1X TBS (20 mM Tris-HCl pH 7.5, 150 mM NaCl). Cell pellets were then resuspended in 700 μL Lysis buffer (50 mM HEPES-KOH pH 7.5, 140 mM NaCl, 1 mM EDTA, 1% Triton X-100, 0.1% Na-deoxycholate and protease inhibitor cocktail (1 mM PMSF, 1 mM Benzamidine, 10 μg/mL Aprotinin, 1 μg/mL Leupeptin, 1 μg/mL Pepstatin)). About the same number of OD_600_ units was used for all samples. Cells were lysed by bead beating and the lysate was sonicated with a Model 100 Sonic dismembrator equipped with a microprobe (Fisher Scientific), 4 x 20 sec at output 7 Watts, with a 1 min break between sonication cycles. Soluble fragmented chromatin was recovered by centrifugation. 600 μL of the chromatin sample was taken per immunoprecipitation (IP) and, when necessary, 6 μL (1%) were saved as an Input sample. The following amounts of antibody per IP were used: anti-Spt16 (a gift from T. Formosa, 1 μL), anti-Pob3 (a gift from T. Formosa, 1 μL), anti-Rpb3 (Neoclone W0012, 3 μL; BioLegend 665004, 1.5 μL), anti-Flag (Sigma-Aldrich F3165, 5 μg), anti-HA (Santa Cruz Biotechnology sc-7392, 15 μL) and rabbit IgG (Millipore 12-370, 2.5 μg). All antibodies have been validated for ChIP (see (Mason and Struhl, 2003) for Spt16 and Pob3 antibodies and manufacturers’ web sites for commercial antibodies). Rabbit antibodies were coupled to Dynabeads coated with Protein G (Thermo Fisher Scientific, 10004D) and mouse antibodies were coupled to Dynabeads coated with Pan Mouse IgG antibodies (Thermo Fisher Scientific, 11042). 50 μL of the appropriate Dynabeads pre-coupled with the indicated antibody were added to the chromatin sample and incubated overnight at 4°C. Beads were washed twice with Lysis buffer, twice with Lysis buffer 500 (Lysis buffer + 360 mM NaCl), twice with Wash buffer (10 mM Tris-HCl pH 8.0, 250 mM LiCl, 0.5% NP40, 0.5% Na-deoxycholate, 1 mM EDTA) and once with TE (10 mM Tris-HCl pH 8.0, 1 mM EDTA). Input and immunoprecipitated chromatin were eluted and reverse-crosslinked with 50 μL TE/SDS (TE + 1% SDS) by incubating overnight at 65°C. Samples were treated with RNase A (345 μL TE, 3 μL 10 mg/mL RNAse A, 2 μL 20 mg/mL Glycogen) at 37°C for 2 hr and subsequently subjected to Proteinase K (15 μL 10% SDS, 7.5 μL 20 mg/mL Proteinase K) digestion at 37°C for 2 hr. Samples were twice phenol/chloroform/isoamyl alcohol (25:24:1) extracted followed by precipitation with 200 mM NaCl and 70% ethanol. Precipitated DNA was resuspended in 50 μL of TE before being used in ChIP-chip experiments.

### ChIP-chip

The ChIP and Input samples (see above) were blunted as follows. 40 µL of ChIP DNA or about 400 ng of Input DNA were added to 70 µL of blunting mix (11 µL 10X NEBuffer 2 (NEB, B7002S), 0.5 µL 10 mg/mL BSA, 0.5 µL 20 mM dNTPs and 0.2 µL (0.6 units) of T4 DNA polymerase (NEB, M0203L)) and incubated at 12°C for 20 min. Then, the blunted DNA was extracted with phenol/chloroform/isoamyl alcohol (25:24:1) in the presence of 280 mM of NaOAc pH 5.2 and 10 μg of glycogen, precipitated with ethanol and resuspended in 25 µL ddH_2_O. Blunted DNA was ligated to 100 pmol of annealed linkers (see **Table S2**) with 2.5 units of T4 DNA ligase (Thermo Fisher Scientific, 15224041) in the supplied buffer, in a final volume of 50 µL, overnight at 16°C. Samples were precipitated with 6 µL of 3 M NaOAc pH 5.2 and 130 µL ethanol, and resuspended in 25 µL ddH_2_O. 15 µL of labeling mix (4 µL 10X ThermoPol Reaction buffer (NEB, B9004S), 2 µL of 5 mM aa-dUTP/dNTP mix (5 mM each dATP, dGTP and dCTP, 3 mM dTTP and 2 mM 5-(3-aminoallyl)-dUTP (Thermo Fisher Scientific, AM8439)) and 1.25 µL of 40 µM oligo 1) were added to each sample. Samples were placed in a thermocycler and when the temperature reached 55°C, 10 µL of enzyme mix (1 µL 10X ThermoPol Reaction buffer, 1 µL (5 units) Taq DNA polymerase (Thermo Fisher Scientific, 18038240) and 0.01 µL (0.025 units) Pfu DNA polymerase (Thermo Fisher Scientific, EP0502)) were added and the following program was run. 1) 72°C, 5 min; 2) 95°C, 2 min; 3) 32 cycles: 95°C, 30sec; 55°C, 30sec; 72°C, 1 min; 4) 72°C, 4 min; 5) 4°C, forever. PCR products were purified using QIAquick PCR Purification kit (QIAGEN, 28106) with few modifications: PE buffer was replaced by Phosphate wash buffer (5 mM KPO_4_ pH 8.5 and 80% ethanol) and EB buffer was replaced by Phosphate elution buffer (4 mM KPO_4_ pH 8.5). Eluates were dried by SpeedVac and pellets were resuspended in 4.5 µL of freshly prepared 0.1 M Na_2_CO_3_ pH 9.0 buffer. 4.5 µL of Cy5 Mono-Reactive NHS Ester Fluorescent Dye (ChIP samples) or Cy3 Mono-Reactive NHS Ester Fluorescent Dye (control samples Input DNA or no-tag ChIP) (GE Healthcare, PA25001 and PA23001) were added. Samples were incubated 1 hr in the dark at room temperature. 35 µL of 0.1 M NaOAc pH 5.2 were added and samples were purified using QIAquick PCR Purification kit according to the manufacturers’ instructions. ChIP and control samples were combined and dried by SpeedVac. Combined samples were resuspended in 110 µL of fresh hybridization mix (5 µL 10 mg/mL Salmon Sperm DNA (Thermo Fisher Scientific, 15632011), 5 µL 8 mg/mL Yeast tRNA (Thermo Fisher Scientific, 15401011) and 100 µL DIG Easy Hyb (Roche, 11603558001)) and incubated 5 min at 95°C followed by 5 min at 45°C. Samples were hybridized to Agilent microarrays in SureHyb enabled chambers (Agilent, G2534A) with rotation at 42°C for at least 18 hr. Microarrays were washed on an orbital shaker for 5 min at room temperature with Wash I buffer (6X SSPE, 0.005% N-Lauroylsarcosine), followed by a second wash with pre-warmed (31°C) Wash II buffer (0.06X SSPE) for 5 min at room temperature. Microarrays were scanned using an InnoScan900 (Innopsys) at 2 µm resolution. The microarrays used were described before (Jeronimo and Robert, 2014) and were custom designed by Agilent Technologies and contain a total of about 180,000 Tm-adjusted 60-mer probes covering the entire yeast genome with virtually no gaps between probes.

### ChIP-exo

ChIP-exo experiments were performed in two biological replicates essentially as described previously (Rossi et al., 2018) with some modifications. After following our standard ChIP protocol using a crosslinked 50 mL cell culture at an OD_600_∼0.8 per IP and the appropriate antibody-coupled Dynabeads (see above), beads were washed twice with Lysis buffer, twice with Lysis buffer 500, twice with Wash buffer and once with 10mM Tris-HCl pH8.0. Beads were then briefly centrifuged to remove all liquid before being used in the ChIP-exo library as follows. For A-tailing of the immunoprecipitated DNA, beads were resuspended in 50 μL of 1X NEBuffer 2 (NEB, B7002S) containing 0.1 mM dATP and 15 units of Klenow Fragment (3’→5’ exo-) (NEB, M0212M) and incubated at 37°C for 30 min. After washing the beads with 10 mM Tris-HCl pH 8.0, dA-tailed DNA was ligated to the First adapter by resuspending the beads in 45 μL of 1X Rapid ligation buffer (Enzymatics, B1010) containing 375 nM of First ligation adapter (synthesized by IDT; see **Table S2** for a list of adapters with their bar codes), 10 units of T4 polynucleotide kinase (NEB, M0201S) and 1,200 units of Rapid T4 DNA ligase (Enzymatics, L6030-HC-L) and incubating at room temperature for 60 min. After washing the beads with 10 mM Tris-HCl pH 8.0, the ligated DNA was subjected to a Fill-in reaction by resuspending the beads in 40 μL of 1X phi29 buffer (NEB) containing 0.2 mg/mL BSA (NEB), 180 μM dNTPs and 10 units of phi29 DNA polymerase (NEB, M0269S) and incubating at 30°C for 20 min. After washing the beads with 10 mM Tris-HCl pH 8.0, DNA was subjected to λ exonuclease digestion by incubating the beads with 50 μL of λ exonuclease reaction buffer (NEB) containing 0.1% Triton X-100, 5% DMSO and 10 units of λ exonuclease (NEB, M0262S) at 37°C for 30 min. To elute and reverse-crosslink DNA, beads were resuspended in 40 uL of reverse crosslink mix containing 25 mM Tris-HCl pH7.5, 200 mM NaCl, 2 mM EDTA, 0.5% SDS and 30 μg of Proteinase K (Thermo Scientific, 25530049) and incubated at 65°C for 16 hr. Eluted DNA was then subjected to a 1.8X cleanup using KAPA Pure Beads (Roche, 07983280001) according to the manufacturers’ instructions before proceeding with Second adapter ligation as follows. Beads were resuspended in 40 μL of 1X T4 DNA ligase reaction buffer (NEB, B0202S) containing 375 nM of Second ligation adapter (synthesized by IDT; see **Table S2**) and 1,200 units of Rapid T4 DNA ligase (Enzymatics, L6030-HC-L), and incubated at room temperature for 60 min. The ligated DNA was then subjected to a 1.8X cleanup with KAPA Pure Beads and eluted in 15 μL ddH_2_O. Libraries were PCR-amplified in 40 μL reactions with 15-18 cycles using KOD Hot Start DNA polymerase (Millipore, 71086-3) and 500 nM of each primer P1.3 and P2.1 (synthesized by IDT; see **Table S2**). At this step, 10 μL from each sample was further amplified for six more cycles to visualize the quality of the library by electrophoresis on a 2% TAE-agarose gel stained with ethidium bromide. Once the library validated, DNA was subjected to a 1X cleanup with KAPA Pure Beads, followed by a double size selection (0.6X-1X) leading to DNA fragments in the 150-450 bp range. Libraries were qualified on Agilent 2100 Bioanalyzer using High Sensitivity DNA kit (Agilent Technologies, 5067-4626) and quantified by qPCR using NEBNext Library Quant kit for Illumina (NEB, E7630). Equal molarity of each library were pooled and subjected to sequencing on an Illumina HiSeq 4000 platform at the McGill University and Génome Québec Innovation Centre to generate 50 bp paired-end reads.

### MNase-ChIP-seq

MNase-ChIP experiments were performed in two biological replicates, as previously described (Rando, 2010) with minor modifications. Briefly, yeast cultures were grown in 500 mL of YPD to an OD_600_ of 0.7-0.9 before crosslinking with 1% formaldehyde (Fisher Scientific, BP531-500) at room temperature for 15 min and quenched with 125 mM glycine at room temperature for 5 min. Crosslinked cells were collected by centrifugation and washed twice with ice-cold ddH_2_O. Yeast cell wall digestion was then performed by resuspending cell pellets in 39 mL Buffer Z (1 M sorbitol and 50 mM Tris-HCl pH 7.4) containing 10 mM β-mercaptoethanol and 10 mg Zymolyase 100T (US Biological, Z1004) and incubating at 30°C with agitation (20-40 min). When digestion was completed, the resulting spheroplasts were pelleted by centrifugation at 3,000 g for 5 min at 4°C and gently resuspended in 2.5 mL of NP buffer (1 M sorbitol, 50 mM NaCl, 10 mM Tris-HCl pH 7.4, 5 mM MgCl_2_, 1 mM CaCl_2_, 500 µM spermidine (Sigma-Aldrich, 85558), 1 mM β-mercaptoethanol and 0.075% NP-40) before being exposed to MNase (Worthington, LS004798) digestion titration to generate the appropriate size of chromatin fragments as follow. 700 µl of spheroplasts were incubated with 4.5, 9, 18, and 36 units of MNase (resuspended in 10 mM Tris-HCl pH 7.4 and 30% glycerol) at 37°C for 20 min and the reaction was stopped by adding 10 mM EDTA. To ensure IP conditions, 200 µL of adjusting buffer (50 mM HEPES-KOH pH7.5, 140 mM NaCl, 1% Triton X-100, and 0.1% Na-deoxycholate) were added to the digested chromatin sample. To determine the appropriate condition of MNase digestion, 45 µL (5%) of each MNase digested sample was taken and incubated with 5 μL of 10X TE/SDS at 65°C for 16 hr to reverse crosslinking. Samples were then treated with RNase A (345 μL TE, 3 μL 10 mg/mL RNAse A, 2 μL 20 mg/mL Glycogen) at 37°C for 2 hr and subsequently subjected to Proteinase K (15 μL 10% SDS, 7.5 μL 20 mg/mL Proteinase K) digestion at 37°C for 2 hr. Samples were twice phenol/chloroform/isoamyl alcohol (25:24:1) extracted followed by precipitation with 200 mM NaCl and 100 % ethanol. Precipitated DNA was resuspended in 30 μL of TE, treated with 1µL of 10 mg/mL RNAse A at 37°C for 1 hr before being analyzed by electrophoresis on a 1.8% TAE-agarose gel stained with ethidium bromide. The digested samples were also qualified on Agilent 2100 Bioanalyzer using High Sensitivity DNA Kit (Agilent Technologies, 5067-4626). The condition giving approximately 65% mono-, 25% di-and 10% tri-nucleosomes was subjected to IP as follows. 45µL (5%) were saved as an Input sample and the remaining digested chromatin was incubated overnight with the appropriate antibody-coupled Dynabeads as per our standard ChIP protocol (see above). Beads were then washed and samples reverse crosslinked, treated with RNase A, and subsequently subjected to Proteinase K before being twice phenol/chloroform/isoamyl alcohol extracted followed by precipitation with NaCl and ethanol. Precipitated DNA was resuspended in 50 μL of TE before being used in sequencing libraries.

Before proceeding with MNase-ChIP-seq libraries, ChIP and Input samples were subjected to a 2X cleanup using KAPA Pure Beads (Roche, 07983280001) according to the manufacturers’ instructions. Samples were eluted with 50 μL of Elution buffer (10 mM Tris pH 8.0) and quantified/qualified on an Agilent 2100 Bioanalyzer instrument using the High Sensitivity DNA kit. 1 ng of Input DNA and 40 μL of ChIP DNA were used for library preparation as follows. The ends of DNA were repaired by incubating in 70 μL of 1X NEBuffer 2 containing 0.6 units of T4 DNA polymerase (NEB, M0203L), 2 units of T4 polynucleotide kinase (NEB, M0201S), 0.09 nM dNTPs and 0.045 μg/μL of BSA at 12°C for 30 min. Repaired DNA was then subjected to a 2X cleanup using KAPA Pure Beads before dA-tailing as follows. Beads containing the repaired DNA were resuspended in 50 μL of 1X NEBuffer 2 containing 0.1 mM dATP and 25 units of Klenow Fragment (3’→5’ exo-) (NEB, M0212M), and incubated at 37°C for 30 min. After a 2X cleanup with KAPA Pure Beads, dA-tailed DNA was ligated to index adapters (Roche, SeqCap Adapter kit A (07141530001) and SeqCap Adapter kit B (07141548001)) as follow. The beads were resuspended in 45 μL of 1X Ligase buffer containing 8 nM of adapter and 2.5 units of T4 DNA ligase and incubated at room temperature for 60 min. The ligated DNA was then subjected to a 1X cleanup with KAPA Pure Beads. Libraries were PCR-amplified with 10-12 cycles using KOD Hot Start DNA polymerase (Millipore, 71086-3) and cleaned up using 1X KAPA Pure Beads. Libraries were qualified on Agilent 2100 Bioanalyzer using High Sensitivity DNA kit and quantified by qPCR using NEBNext Library Quant kit for Illumina (NEB, E7630). Equal molarity of each library were pooled and subjected to sequencing on Illumina HiSeq 4000 and NovaSeq 6000 platforms at the McGill University and Génome Québec Innovation Centre to generate 50 bp paired-end reads.

### Co-immunoprecipitation experiments

For co-IP experiments between Spt16 and Chd1 in WT and *chd1*-K407R cells, yeast strains were grown in 50 mL of YPD medium to an OD_600_ of 0.7-0.9. Cells were collected by centrifugation, washed in cold ddH_2_O, and resuspended in 700 μL of IP buffer (50 mM HEPES-KOH pH 7.5, 100 mM NaCl, 0.5 mM EDTA, 0.05% NP-40, 20% glycerol, 1 mM DTT and (1 mM PMSF, 1 mM Benzamidine, 10 μg/mL Aprotinin, 1 μg/mL Leupeptin, 1 μg/mL Pepstatin)). About the same amount of OD_600_ units was used for all samples. Cells were lysed by bead beating for 5 min and the soluble extract was recovered by centrifugation (15 min at 4°C at 18,000 × g). About 3-3.5 mg of protein (600 µl of cleared lysate) was taken per IP and 6 μL (1%) were saved as Input sample. The cleared lysate was incubated for 60 min at 4°C with 50 μL of pre-washed magnetic Pan Mouse Dynabeads (Thermo Fisher Scientific) coupled to 15 μL of anti-HA F7 mouse monoclonal antibody (Santa Cruz Biotechnology). Samples were then washed three times with IP buffer and resuspended in Laemmli buffer. Samples were incubated at 95^°^C for 5 min, briefly centrifuged and analyzed by SDS-PAGE and Western blot using anti-Spt16 rabbit polyclonal (a gift from T. Formosa, 1:1000 dilution) and anti-HA mouse monoclonal (Santa Cruz Biotechnology sc-7392, 1:1000 dilution) antibodies. Membranes were incubated with donkey anti-rabbit IRDye 800CW and donkey anti-mouse IRDye 680RD antibodies (LI-COR Biosciences, 926-32213 and 926-68072) according to the manufacturers’ instructions and scanned on the Odyssey infrared imaging system (LI-COR Biosciences).

## QUANTIFICATION AND STATISTICAL ANALYSIS

### ChIP-chip Data analysis

The ChIP-chip data were normalized using the Limma Loess method, and replicates were combined as described previously (Ren et al., 2000). The data were subjected to one round of smoothing using a Gaussian sliding window with a standard deviation of 100 bp to generate data points in 10 bp intervals (Guillemette et al., 2005).

### ChIP-exo Data analysis

Paired-end reads from ChIP-exo data were aligned to the yeast *S*. *cerevisiae* genome (sacCer3 from UCSC Genome Browser) using BWA version 0.7.17 (Li and Durbin, 2009). Unmapped reads, reads not mapped properly in pairs, and duplicates were removed from the analysis using SAMtools version 1.8 (Li et al., 2009). For each paired-end read, only the first read was kept for further analysis. The aligned files were converted to BED using BEDTools version 2.28.0 (Quinlan and Hall, 2010). BED files containing reads from replicates were further analyzed individually in addition to being merged. The reads were moved 6 bp towards the 3’ end to correct for the offset of exonuclease stops and the site of crosslinking using a custom java script called bed-tools-j version 2.1 available at https://github.com/francoisrobertlab/bed-tools-j/releases/tag/2.1 with the parameter “moveannotations -d 6 -r -dn”. Three genome coverage files were computed using the 5’ position of the moved reads with BEDTools. Coverage files were generated for reads mapping the positive strand, the negative strand, and the merged strands. The genome coverage files were converted to the bigWig format using the utilities from UCSC Genome Browser (Casper et al., 2018). The analysis was partially automated using a custom script called seqtools version 1.0 available at https://github.com/francoisrobertlab/seqtools.

### MNase-ChIP-seq Data analysis

Paired-end reads from MNase-ChIP-seq data where aligned on the *S*. *cerevisiae* genome version sacCer3 using Bowtie2 version 2.3.4.3 (Langmead and Salzberg, 2012) with the parameter “-X 1000” to include longer fragments. Unmapped and low-quality fragments were removed using SAMtools version 1.9 (Li et al., 2009) with the following parameter “view -f 2 -F 2048”. Duplicates were removed using SAMtools with the following parameters “fixmate -m” and “markdup -r”. The aligned files were converted to BED with BEDTools version 2.27.1 (Quinlan and Hall, 2010) using seqtools with the command “seqtools bam2bed” resulting in BED files containing the DNA fragments. The BED files of replicates were merged using seqtools with command “seqtools merge”. Genome coverage files were generated using seqtools with commands “mnasetools prepgenomecov” and “seqtools genomecov” that require BEDTools version 2.27.1 and KentUtils (downloaded on July 16^th^ 2018). The genome coverage files were converted to the bigWig format using the utilities from UCSC Genome Browser (Casper et al., 2018). The configuration files used with seqtools for the MNase-ChIP-seq data analysis are available on GitHub at https://github.com/francoisrobertlab/mnasechip-cj2020.

### Heatmaps of coverage over genes versus fragment size

To generate **Figures 3B, S3B** and **S3D**, fragments from the MNase-ChIP-seq data were grouped in bins of 10 bp using seqtools with command “seqtools split --binLength 10 --binMinLength 50 --binMaxLength 500” and genome coverage files were generated using seqtools with commands “mnasetools prepgenomecov” and “seqtools genomecov” that require BEDTools version 2.27.1 (Quinlan and Hall, 2010) and KentUtils (downloaded on July 16^th^ 2018). The heatmap matrices were then generated using seqtools with command “seqtools vap” that requires VAP (Coulombe et al., 2014) (Note: split and genome coverage must be executed prior to “seqtools vap”). Heatmaps were then generated using Java TreeView Version 1.1.6r4 (Saldanha, 2004).

### Two-dimensional occupancy (2DO) plots

In **Figures 4** and **S4**, Plot2DO (Beati and Chereji, 2020) revision number 87fadb4acd23214f83e5440b0ccb02c623fa62d9 with parameters “-t dyads -r Plus1 -l 100 -L 400 -m 0.02” was used to generate 2DO heatmaps from the MNase-ChIP-seq data. In some analyzes, the fragments not overlapping the top 10% of genes were removed using seqtools with command “seqtools intersect -a Top10percentTxbdGenes.txt”, where “Top10percentTxbdGenes.txt” contains the top 10% most transcribed genes and their coordinates.

### Identification of coordinates of +2 nucleosomes

Coordinates of +1 nucleosomes were from (Chereji et al., 2018) while coordinates of +2 nucleosomes were determined from the chemical cleavage data (Chereji et al., 2018) as follow. BigWig dyads from (Chereji et al., 2018) were downloaded and merged using seqtools with command “seqtools mergebw”. Then the position of the +2 dyad was computed using seqtools with command “seqtools dyadposition -g 13059_2018_1398_MOESM2_ESM.txt -s merge.bw”, where “13059_2018_1398_MOESM2_ESM.txt” is the file from (Chereji et al., 2018) containing the +1 nucleosomes coordinates. The seqtools script assesses the position of the +2 nucleosome by finding the position of the maximum signal from the merged bigWig file at a distance between 141 and 191 bp from the position of the +1 nucleosome for each gene.

### Distributions of MNase-ChIP-seq fragments relative to nucleosome dyads

In **Figures 6B** and **S6**, the distribution of the center of MNase-ChIP-seq DNA fragments of different size, relative to the dyad of the +1 or +2 nucleosomes were generated as follow. The fragments from MNase-ChIP-seq data were grouped in bins of 10 bp using seqtools with commands “seqtools slowsplit -- binMinLength 63 --binMaxLength 73”, “seqtools slowsplit --binMinLength 85 --binMaxLength 95”, “seqtools slowsplit --binMinLength 98 --binMaxLength 108” and, “seqtools slowsplit --binMinLength 120 --binMaxLength 130”. The genome coverage was computed from the resulting files using seqtools with command “mnasetools prepgenomecov” and then “seqtools genomecov --scale 1.0” which requires BEDTools 2.27.1 (Quinlan and Hall, 2010) and KentUtils (downloaded on July 16th 2018). The dyad coverage was computed from the genome coverage files using seqtools with command “mnasetools dyadcov -g first_dyad.txt --smoothing 20”, where “first_dyad.txt” contains the position of the +1 nucleosome from (Chereji et al., 2018). The dyad coverage for the second dyad used “mnasetools dyadcov -g second_dyad.txt --smoothing 20”, where “second_dyad.txt” contains the position of the +2 nucleosome from “seqtools dyadposition” execution. Gaussian fits were computed using seqtools with command “seqtools fitgaussian--svg --amin 0”, while double Gaussian fits were computed using seqtools with command “seqtools fitdoublegaussian --gaussian --svg --amin1 0 --amin2 0”. The fitting was computed using the lmfit Python library version 1.0.0 available at https://lmfit.github.io/lmfit-py/ using the GaussianModel and ConstantModel.

### Mathematical modeling

Models were implemented in Matlab 2019b and used a fixed time-step stochastic algorithm in which it is assumed that only one reaction can occur in each time step. At each time step, a random number was drawn from a uniform distribution (Matlab rand() function, Mersenne Twister random number generator). This random number was then compared against a list of the cumulative summed probabilities per unit time of all possible reactions as described in the pseudo-code below (with description in roman font and pseudo code in italics). Table 1 contains parameter values and their abbreviations as used in the pseudo-code. The simulation keeps track of the number of FACT complexes and the numbers that are in the extending, retracting, fully extended and fully retracted states, FACTNo, ExtNo, RetNo, FullyExtNo and FullyRetNo, respectively.

Generate random number:

*Diceroll = rand;*

Compare random number against possible reactions:

*if rand < (BR)*

Bind a FACT complex at the +1 Nucleosome location (if it is unoccupied)

*else if rand < (BR+ExtNo*ER)*

Select a random FACT complex from those that are in the extending state and, if possible, extend by one base pair at the TSS-distal side – if the complex abuts with a preceding complex, do not extend; if the complexes leading edge reaches the end of the gene, flag as fully extended

*else if rand < (BR+ExtNo*ER+RetNo*RR)*

Select a random FACT complex from those that are in the retracting state and, if possible, retract by one base pair at the TSS-proximal side

*else if rand < (BR+ExtNo*ER+RetNo*RR+FullyExtNo*ERS)*

Select a random FACT complex from those that are in the fully extended state and transition to the retracting state

*else if rand < (BR+ExtNo*ER+RetNo*RR+FullyExtNo*ERS+FullyRetNo*RES)*

Select a random FACT complex from those that are in the fully retracted state and transition to the extending state – if the FACT complex has reached the end of the gene, unbind

*else if rand < (BR+ExtNo*ER+RetNo*RR+FullyExtNo*ERS+FullyRetNo*RES+FactNo*UBR)* Select a random FACT complex and unbind

*else*

Leave the system unchanged

*end*

The system was simulated on a synthetic gene of length 1,510 bp. Nucleosome locations were defined as regions of 150 bp, with the first at the beginning of the gene and spacing regions of 20 bp between each nucleosome location. The minimum extent of a FACT complex was taken to be 150 bp and the maximum to be 320 bp (the extent of two nucleosomes and a spacing region).

The system was simulated for a relaxation period of 10^6^ time steps, following this the simulation was continued for an additional 10^6^ during which a measurement of protected fragment sizes and positions was taken every 10-time steps. Where presented in figures, the fragment sizes and positions are displayed as block averages – each displayed pixel is the mean of the 20-by-20 region around it in the original measurement. This was to increase visibility as the simulations lack noise in the positioning and size of nucleosomes and FACT complexes.

In addition to the core mechanisms described above, the possibility for new FACT complexes to bind with uniform probability to any unoccupied nucleosome location in the gene was tested.

**Table 1:** Parameter values, expressed as probabilities per unit time, and their abbreviations used in the main model for the WT and *chd1*Δ cases (which differ only in the extension rate), and the internal binding model, see **Figure 5**.

| Parameter | Abbr. | WT (Fig. 5B) | *chd1Δ* (Figs. 5D, 5G) | Internal Binding (Fig. 5C) |
| --- | --- | --- | --- | --- |
| Binding | *BR* | 0.00005 | 0.00005 | 0.00045 |
| Extension | *ER* | 0.03 | 0.0075 | 0.03 |
| Extension to Retraction | *ERS* | 0.003 | 0.003 | 0.003 |
| Retraction | *RR* | 0.03 | 0.03 | 0.03 |
| Retraction to Extension | *RES* | 0.003 | 0.003 | 0.003 |
| Unbinding | *UBR* | 0.000005 | 0.000005 | 0 |
| Relaxation Time Steps | TSR | $10^6$ | $10^6$ | $10^6$ |
| Measurement Time Steps | TSM | $10^6$ | $10^6$ | $10^6$ |
| Measurement Interval | MI | 10 | 10 | 10 |
| Realisations | R | 10000 | 10000 | 10000 |

**Table 2:**
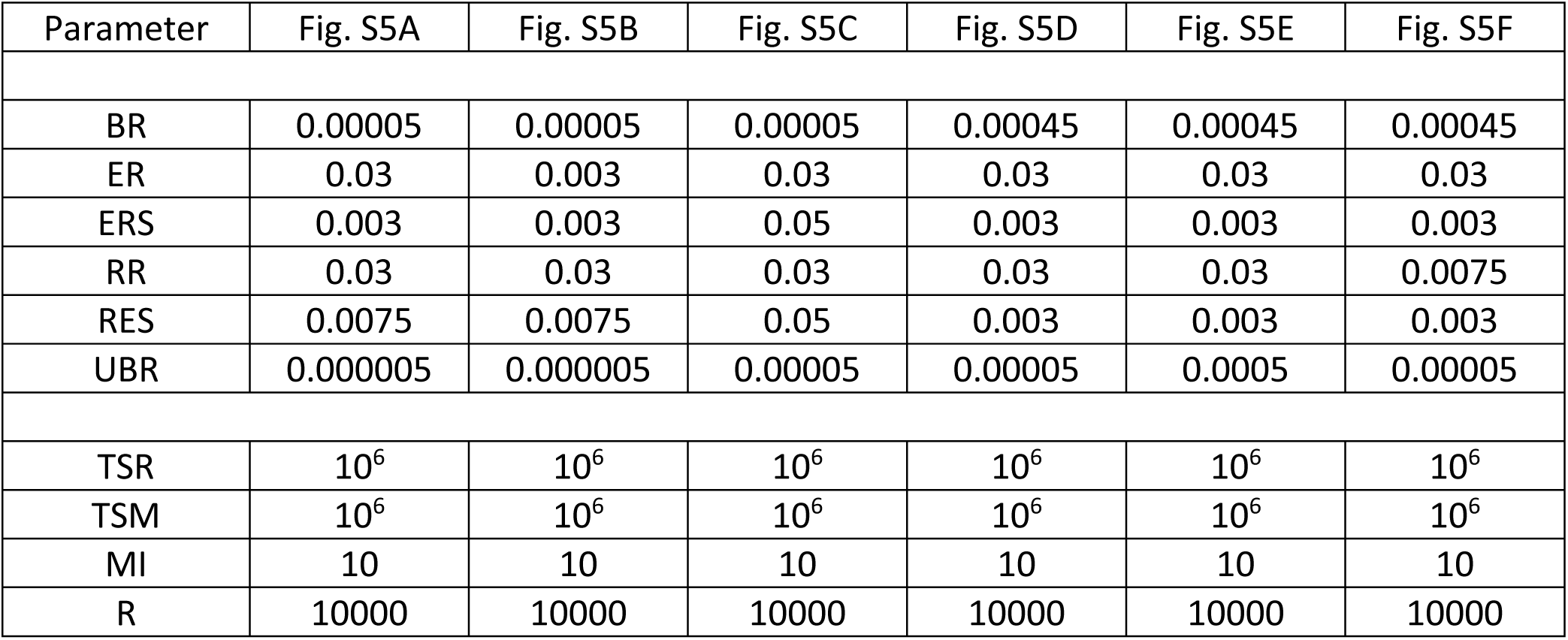
Parameter values, expressed as probabilities per unit time used for the modelling data shown in **Figure S5**.

| Parameter | Fig. S5A | Fig. S5B | Fig. S5C | Fig. S5D | Fig. S5E | Fig. S5F |
| --- | --- | --- | --- | --- | --- | --- |
| BR | 0.00005 | 0.00005 | 0.00005 | 0.00045 | 0.00045 | 0.00045 |
| ER | 0.03 | 0.003 | 0.03 | 0.03 | 0.03 | 0.03 |
| ERS | 0.003 | 0.003 | 0.05 | 0.003 | 0.003 | 0.003 |
| RR | 0.03 | 0.03 | 0.03 | 0.03 | 0.03 | 0.0075 |
| RES | 0.0075 | 0.0075 | 0.05 | 0.003 | 0.003 | 0.003 |
| UBR | 0.000005 | 0.000005 | 0.00005 | 0.00005 | 0.0005 | 0.00005 |
| TSR | $10^6$ | $10^6$ | $10^6$ | $10^6$ | $10^6$ | $10^6$ |
| TSM | $10^6$ | $10^6$ | $10^6$ | $10^6$ | $10^6$ | $10^6$ |
| MI | 10 | 10 | 10 | 10 | 10 | 10 |
| R | 10000 | 10000 | 10000 | 10000 | 10000 | 10000 |

## KEY RESOURCE TABLE

See accompanying document.

## Notes

### Competing Interest Statement

The authors have declared no competing interest.

## REFERENCES

Adelman, K., Wei, W., Ardehali, M.B., Werner, J., Zhu, B., Reinberg, D., and Lis, J.T. (2006). Drosophila Paf1 modulates chromatin structure at actively transcribed genes. Mol Cell Biol 26, 250–260.

Beati, P., and Chereji, R.V. (2020). Creating 2D Occupancy Plots Using plot2DO. Methods Mol Biol 2117, 93–108.

Belotserkovskaya, R., Oh, S., Bondarenko, V.A., Orphanides, G., Studitsky, V.M., and Reinberg, D. (2003). FACT facilitates transcription-dependent nucleosome alteration. Science 301, 1090–1093.

Birch, J.L., Tan, B.C., Panov, K.I., Panova, T.B., Andersen, J.S., Owen-Hughes, T.A., Russell, J., Lee, S.C., and Zomerdijk, J.C. (2009). FACT facilitates chromatin transcription by RNA polymerases I and III. EMBO J 28, 854–865.

Biswas, D., Dutta-Biswas, R., Mitra, D., Shibata, Y., Strahl, B.D., Formosa, T., and Stillman, D.J. (2006). Opposing roles for Set2 and yFACT in regulating TBP binding at promoters. EMBO J 25, 4479–4489.

Biswas, D., Yu, Y., Prall, M., Formosa, T., and Stillman, D.J. (2005). The yeast FACT complex has a role in transcriptional initiation. Mol Cell Biol 25, 5812–5822.

Carrozza, M.J., Li, B., Florens, L., Suganuma, T., Swanson, S.K., Lee, K.K., Shia, W.J., Anderson, S., Yates, J., Washburn, M.P., et al. (2005). Histone H3 methylation by Set2 directs deacetylation of coding regions by Rpd3S to suppress spurious intragenic transcription. Cell 123, 581–592.

Carvalho, S., Raposo, A.C., Martins, F.B., Grosso, A.R., Sridhara, S.C., Rino, J., Carmo-Fonseca, M., and de Almeida, S.F. (2013). Histone methyltransferase SETD2 coordinates FACT recruitment with nucleosome dynamics during transcription. Nucleic Acids Res 41, 2881–2893.

Casper, J., Zweig, A.S., Villarreal, C., Tyner, C., Speir, M.L., Rosenbloom, K.R., Raney, B.J., Lee, C.M., Lee, B.T., Karolchik, D., et al. (2018). The UCSC Genome Browser database: 2018 update. Nucleic Acids Res 46, D762–D769.

Chereji, R.V., Ramachandran, S., Bryson, T.D., and Henikoff, S. (2018). Precise genome-wide mapping of single nucleosomes and linkers in vivo. Genome Biol 19, 19.

Collin, P., Jeronimo, C., Poitras, C., and Robert, F. (2019). RNA Polymerase II CTD Tyrosine 1 Is Required for Efficient Termination by the Nrd1-Nab3-Sen1 Pathway. Mol Cell 73, 655–669 e657.

Coulombe, C., Poitras, C., Nordell-Markovits, A., Brunelle, M., Lavoie, M.A., Robert, F., and Jacques, P.E. (2014). VAP: a versatile aggregate profiler for efficient genome-wide data representation and discovery. Nucleic Acids Res 42, W485–493.

Crickard, J.B., Lee, J., Lee, T.H., and Reese, J.C. (2017). The elongation factor Spt4/5 regulates RNA polymerase II transcription through the nucleosome. Nucleic Acids Res 45, 6362–6374.

Doris, S.M., Chuang, J., Viktorovskaya, O., Murawska, M., Spatt, D., Churchman, L.S., and Winston, F. (2018). Spt6 Is Required for the Fidelity of Promoter Selection. Mol Cell 72, 687–699 e686.

Drouin, S., Laramee, L., Jacques, P.E., Forest, A., Bergeron, M., and Robert, F. (2010). DSIF and RNA polymerase II CTD phosphorylation coordinate the recruitment of Rpd3S to actively transcribed genes. PLoS Genet 6, e1001173.

Ehara, H., Kujirai, T., Fujino, Y., Shirouzu, M., Kurumizaka, H., and Sekine, S.I. (2019). Structural insight into nucleosome transcription by RNA polymerase II with elongation factors. Science 363, 744–747.

Ehara, H., Yokoyama, T., Shigematsu, H., Yokoyama, S., Shirouzu, M., and Sekine, S.I. (2017). Structure of the complete elongation complex of RNA polymerase II with basal factors. Science 357, 921–924.

Farnung, L., Vos, S.M., and Cramer, P. (2018). Structure of transcribing RNA polymerase II-nucleosome complex. Nat Commun 9, 5432.

Farnung, L., Vos, S.M., Wigge, C., and Cramer, P. (2017). Nucleosome-Chd1 structure and implications for chromatin remodelling. Nature 550, 539–542.

Fleming, A.B., Kao, C.F., Hillyer, C., Pikaart, M., and Osley, M.A. (2008). H2B ubiquitylation plays a role in nucleosome dynamics during transcription elongation. Mol Cell 31, 57–66.

Foltman, M., Evrin, C., De Piccoli, G., Jones, R.C., Edmondson, R.D., Katou, Y., Nakato, R., Shirahige, K., and Labib, K. (2013). Eukaryotic replisome components cooperate to process histones during chromosome replication. Cell Rep 3, 892–904.

Formosa, T. (2013). The role of FACT in making and breaking nucleosomes. Biochim Biophys Acta 1819, 247–255.

Formosa, T., Eriksson, P., Wittmeyer, J., Ginn, J., Yu, Y., and Stillman, D.J. (2001). Spt16-Pob3 and the HMG protein Nhp6 combine to form the nucleosome-binding factor SPN. EMBO J 20, 3506–3517.

Govind, C.K., Qiu, H., Ginsburg, D.S., Ruan, C., Hofmeyer, K., Hu, C., Swaminathan, V., Workman, J.L., Li, B., and Hinnebusch, A.G. (2010). Phosphorylated Pol II CTD recruits multiple HDACs, including Rpd3C(S), for methylation-dependent deacetylation of ORF nucleosomes. Mol Cell 39, 234–246.

Guillemette, B., Bataille, A.R., Gevry, N., Adam, M., Blanchette, M., Robert, F., and Gaudreau, L. (2005). Variant histone H2A.Z is globally localized to the promoters of inactive yeast genes and regulates nucleosome positioning. PLoS Biol 3, e384.

Gurova, K., Chang, H.W., Valieva, M.E., Sandlesh, P., and Studitsky, V.M. (2018). Structure and function of the histone chaperone FACT - Resolving FACTual issues. Biochim Biophys Acta Gene Regul Mech 1861, 892–904.

Han, J., Li, Q., McCullough, L., Kettelkamp, C., Formosa, T., and Zhang, Z. (2010). Ubiquitylation of FACT by the cullin-E3 ligase Rtt101 connects FACT to DNA replication. Genes Dev 24, 1485–1490.

Henikoff, J.G., Belsky, J.A., Krassovsky, K., MacAlpine, D.M., and Henikoff, S. (2011). Epigenome characterization at single base-pair resolution. Proc Natl Acad Sci U S A 108, 18318–18323.

Jamai, A., Puglisi, A., and Strubin, M. (2009). Histone chaperone spt16 promotes redeposition of the original h3-h4 histones evicted by elongating RNA polymerase. Mol Cell 35, 377–383.

Jeronimo, C., Poitras, C., and Robert, F. (2019). Histone Recycling by FACT and Spt6 during Transcription Prevents the Scrambling of Histone Modifications. Cell Rep 28, 1206–1218 e1208.

Jeronimo, C., and Robert, F. (2014). Kin28 regulates the transient association of Mediator with core promoters. Nat Struct Mol Biol 21, 449–455.

Jeronimo, C., Watanabe, S., Kaplan, C.D., Peterson, C.L., and Robert, F. (2015). The Histone Chaperones FACT and Spt6 Restrict H2A.Z from Intragenic Locations. Mol Cell 58, 1113–1123.

Jiang, C., and Pugh, B.F. (2009). A compiled and systematic reference map of nucleosome positions across the Saccharomyces cerevisiae genome. Genome Biol 10, R109.

John, S., Howe, L., Tafrov, S.T., Grant, P.A., Sternglanz, R., and Workman, J.L. (2000). The something about silencing protein, Sas3, is the catalytic subunit of NuA3, a yTAF(II)30-containing HAT complex that interacts with the Spt16 subunit of the yeast CP (Cdc68/Pob3)-FACT complex. Genes Dev 14, 1196–1208.

Jonkers, I., and Lis, J.T. (2015). Getting up to speed with transcription elongation by RNA polymerase II. Nat Rev Mol Cell Biol 16, 167–177.

Krogan, N.J., Cagney, G., Yu, H., Zhong, G., Guo, X., Ignatchenko, A., Li, J., Pu, S., Datta, N., Tikuisis, A.P., et al. (2006). Global landscape of protein complexes in the yeast Saccharomyces cerevisiae. Nature 440, 637–643.

Krogan, N.J., Kim, M., Ahn, S.H., Zhong, G., Kobor, M.S., Cagney, G., Emili, A., Shilatifard, A., Buratowski, S., and Greenblatt, J.F. (2002). RNA polymerase II elongation factors of Saccharomyces cerevisiae: a targeted proteomics approach. Mol Cell Biol 22, 6979–6992.

Krogan, N.J., Kim, M., Tong, A., Golshani, A., Cagney, G., Canadien, V., Richards, D.P., Beattie, B.K., Emili, A., Boone, C., et al. (2003). Methylation of histone H3 by Set2 in Saccharomyces cerevisiae is linked to transcriptional elongation by RNA polymerase II. Mol Cell Biol 23, 4207–4218.

Krogan, N.J., Peng, W.T., Cagney, G., Robinson, M.D., Haw, R., Zhong, G., Guo, X., Zhang, X., Canadien, V., Richards, D.P., et al. (2004). High-definition macromolecular composition of yeast RNA-processing complexes. Mol Cell 13, 225–239.

Kujirai, T., Ehara, H., Fujino, Y., Shirouzu, M., Sekine, S.I., and Kurumizaka, H. (2018). Structural basis of the nucleosome transition during RNA polymerase II passage. Science 362, 595–598.

Kwon, S.H., Florens, L., Swanson, S.K., Washburn, M.P., Abmayr, S.M., and Workman, J.L. (2010). Heterochromatin protein 1 (HP1) connects the FACT histone chaperone complex to the phosphorylated CTD of RNA polymerase II. Genes Dev 24, 2133–2145.

Lai, W.K.M., and Pugh, B.F. (2017). Understanding nucleosome dynamics and their links to gene expression and DNA replication. Nat Rev Mol Cell Biol 18, 548–562.

Langmead, B., and Salzberg, S.L. (2012). Fast gapped-read alignment with Bowtie 2. Nat Methods 9, 357–359.

Laughery, M.F., Hunter, T., Brown, A., Hoopes, J., Ostbye, T., Shumaker, T., and Wyrick, J.J. (2015). New vectors for simple and streamlined CRISPR-Cas9 genome editing in Saccharomyces cerevisiae. Yeast 32, 711–720.

Li, H., and Durbin, R. (2009). Fast and accurate short read alignment with Burrows-Wheeler transform. Bioinformatics 25, 1754–1760.

Li, H., Handsaker, B., Wysoker, A., Fennell, T., Ruan, J., Homer, N., Marth, G., Abecasis, G., Durbin, R., and Genome Project Data Processing, S. (2009). The Sequence Alignment/Map format and SAMtools. Bioinformatics 25, 2078–2079.

Lindstrom, D.L., Squazzo, S.L., Muster, N., Burckin, T.A., Wachter, K.C., Emigh, C.A., McCleery, J.A., Yates, J.R., 3rd, and Hartzog, G.A. (2003). Dual roles for Spt5 in pre-mRNA processing and transcription elongation revealed by identification of Spt5-associated proteins. Mol Cell Biol 23, 1368–1378.

Liu, Y., Zhou, K., Zhang, N., Wei, H., Tan, Y.Z., Zhang, Z., Carragher, B., Potter, C.S., D’Arcy, S., and Luger, K. (2020). FACT caught in the act of manipulating the nucleosome. Nature 577, 426–431.

Martin, B.J.E., Chruscicki, A.T., and Howe, L.J. (2018). Transcription Promotes the Interaction of the FAcilitates Chromatin Transactions (FACT) Complex with Nucleosomes in Saccharomyces cerevisiae. Genetics 210, 869–881.

Mason, P.B., and Struhl, K. (2003). The FACT complex travels with elongating RNA polymerase II and is important for the fidelity of transcriptional initiation in vivo. Mol Cell Biol 23, 8323–8333.

McCullough, L.L., Connell, Z., Xin, H., Studitsky, V.M., Feofanov, A.V., Valieva, M.E., and Formosa, T. (2018). Functional roles of the DNA-binding HMGB domain in the histone chaperone FACT in nucleosome reorganization. J Biol Chem 293, 6121–6133.

Nesher, E., Safina, A., Aljahdali, I., Portwood, S., Wang, E.S., Koman, I., Wang, J., and Gurova, K.V. (2018). Role of Chromatin Damage and Chromatin Trapping of FACT in Mediating the Anticancer Cytotoxicity of DNA-Binding Small-Molecule Drugs. Cancer Res 78, 1431–1443.

Orphanides, G., LeRoy, G., Chang, C.H., Luse, D.S., and Reinberg, D. (1998). FACT, a factor that facilitates transcript elongation through nucleosomes. Cell 92, 105–116.

Pathak, R., Singh, P., Ananthakrishnan, S., Adamczyk, S., Schimmel, O., and Govind, C.K. (2018). Acetylation-Dependent Recruitment of the FACT Complex and Its Role in Regulating Pol II Occupancy Genome-Wide in Saccharomyces cerevisiae. Genetics 209, 743–756.

Petrenko, N., Jin, Y., Dong, L., Wong, K.H., and Struhl, K. (2019). Requirements for RNA polymerase II preinitiation complex formation in vivo. eLife 8, e43654.

Quinlan, A.R., and Hall, I.M. (2010). BEDTools: a flexible suite of utilities for comparing genomic features. Bioinformatics 26, 841–842.

Ramachandran, S., Ahmad, K., and Henikoff, S. (2017). Transcription and Remodeling Produce Asymmetrically Unwrapped Nucleosomal Intermediates. Mol Cell 68, 1038–1053 e1034.

Rando, O.J. (2010). Genome-wide mapping of nucleosomes in yeast. Methods Enzymol 470, 105–118.

Ren, B., Robert, F., Wyrick, J.J., Aparicio, O., Jennings, E.G., Simon, I., Zeitlinger, J., Schreiber, J., Hannett, N., Kanin, E., et al. (2000). Genome-wide location and function of DNA binding proteins. Science 290, 2306–2309.

Rossi, M.J., Lai, W.K.M., and Pugh, B.F. (2018). Simplified ChIP-exo assays. Nat Commun 9, 2842.

Ruone, S., Rhoades, A.R., and Formosa, T. (2003). Multiple Nhp6 molecules are required to recruit Spt16-Pob3 to form yFACT complexes and to reorganize nucleosomes. J Biol Chem 278, 45288–45295.

Saldanha, A.J. (2004). Java Treeview--extensible visualization of microarray data. Bioinformatics 20, 3246–3248.

Schneider, B.L., Seufert, W., Steiner, B., Yang, Q.H., and Futcher, A.B. (1995). Use of polymerase chain reaction epitope tagging for protein tagging in Saccharomyces cerevisiae. Yeast 11, 1265–1274.

Sdano, M.A., Fulcher, J.M., Palani, S., Chandrasekharan, M.B., Parnell, T.J., Whitby, F.G., Formosa, T., and Hill, C.P. (2017). A novel SH2 recognition mechanism recruits Spt6 to the doubly phosphorylated RNA polymerase II linker at sites of transcription. eLife 6, e28723.

Sen, R., Kaja, A., Ferdoush, J., Lahudkar, S., Barman, P., and Bhaumik, S.R. (2017). An mRNA Capping Enzyme Targets FACT to the Active Gene To Enhance the Engagement of RNA Polymerase II into Transcriptional Elongation. Mol Cell Biol 37, e00029–00017.

Simic, R., Lindstrom, D.L., Tran, H.G., Roinick, K.L., Costa, P.J., Johnson, A.D., Hartzog, G.A., and Arndt, K.M. (2003). Chromatin remodeling protein Chd1 interacts with transcription elongation factors and localizes to transcribed genes. EMBO J 22, 1846–1856.

Skene, P.J., Hernandez, A.E., Groudine, M., and Henikoff, S. (2014). The nucleosomal barrier to promoter escape by RNA polymerase II is overcome by the chromatin remodeler Chd1. eLife 3, e02042.

Smart, S.K., Mackintosh, S.G., Edmondson, R.D., Taverna, S.D., and Tackett, A.J. (2009). Mapping the local protein interactome of the NuA3 histone acetyltransferase. Protein Sci 18, 1987–1997.

Sun, X.M., Bowman, A., Priestman, M., Bertaux, F., Martinez-Segura, A., Tang, W., Whilding, C., Dormann, D., Shahrezaei, V., and Marguerat, S. (2020). Size-Dependent Increase in RNA Polymerase II Initiation Rates Mediates Gene Expression Scaling with Cell Size. Curr Biol 30, 1217–1230 e1217.

Sundaramoorthy, R., Hughes, A.L., El-Mkami, H., Norman, D.G., Ferreira, H., and Owen-Hughes, T. (2018). Structure of the chromatin remodelling enzyme Chd1 bound to a ubiquitinylated nucleosome. eLife 7, e35720.

Takahata, S., Yu, Y., and Stillman, D.J. (2009). FACT and Asf1 regulate nucleosome dynamics and coactivator binding at the HO promoter. Mol Cell 34, 405–415.

Tessarz, P., Santos-Rosa, H., Robson, S.C., Sylvestersen, K.B., Nelson, C.J., Nielsen, M.L., and Kouzarides, T. (2014). Glutamine methylation in histone H2A is an RNA-polymerase-I-dedicated modification. Nature 505, 564–568.

Teves, S.S., Weber, C.M., and Henikoff, S. (2014). Transcribing through the nucleosome. Trends Biochem Sci 39, 577–586.

Tsunaka, Y., Fujiwara, Y., Oyama, T., Hirose, S., and Morikawa, K. (2016). Integrated molecular mechanism directing nucleosome reorganization by human FACT. Genes Dev 30, 673–686.

Venkatesh, S., and Workman, J.L. (2015). Histone exchange, chromatin structure and the regulation of transcription. Nat Rev Mol Cell Biol 16, 178–189.

Vinayachandran, V., Reja, R., Rossi, M.J., Park, B., Rieber, L., Mittal, C., Mahony, S., and Pugh, B.F. (2018). Widespread and precise reprogramming of yeast protein-genome interactions in response to heat shock. Genome Res 28, 357–366.

Vos, S.M., Farnung, L., Boehning, M., Wigge, C., Linden, A., Urlaub, H., and Cramer, P. (2018). Structure of activated transcription complex Pol II-DSIF-PAF-SPT6. Nature 560, 607–612.

Vos, S.M., Farnung, L., Linden, A., Urlaub, H., and Cramer, P. (2020). Structure of complete Pol II-DSIF-PAF-SPT6 transcription complex reveals RTF1 allosteric activation. Nat Struct Mol Biol 27, 668–677.

Voth, W.P., Takahata, S., Nishikawa, J.L., Metcalfe, B.M., Naar, A.M., and Stillman, D.J. (2014). A role for FACT in repopulation of nucleosomes at inducible genes. PLoS One 9, e84092.

Wang, T., Liu, Y., Edwards, G., Krzizike, D., Scherman, H., and Luger, K. (2018). The histone chaperone FACT modulates nucleosome structure by tethering its components. Life Sci Alliance 1, e201800107.

Wong, K.H., Jin, Y., and Struhl, K. (2014). TFIIH phosphorylation of the Pol II CTD stimulates mediator dissociation from the preinitiation complex and promoter escape. Mol Cell 54, 601–612.

Xu, Y., Bernecky, C., Lee, C.T., Maier, K.C., Schwalb, B., Tegunov, D., Plitzko, J.M., Urlaub, H., and Cramer, P. (2017). Architecture of the RNA polymerase II-Paf1C-TFIIS transcription elongation complex. Nat Commun 8, 15741.

Zhou, K., Liu, Y., and Luger, K. (2020). Histone chaperone FACT FAcilitates Chromatin Transcription: mechanistic and structural insights. Curr Opin Struct Biol 65, 26–32.

