## Supplemental material for "FACT is recruited to the +1 nucleosome of transcribed genes and spreads in a Chd1-dependent manner"

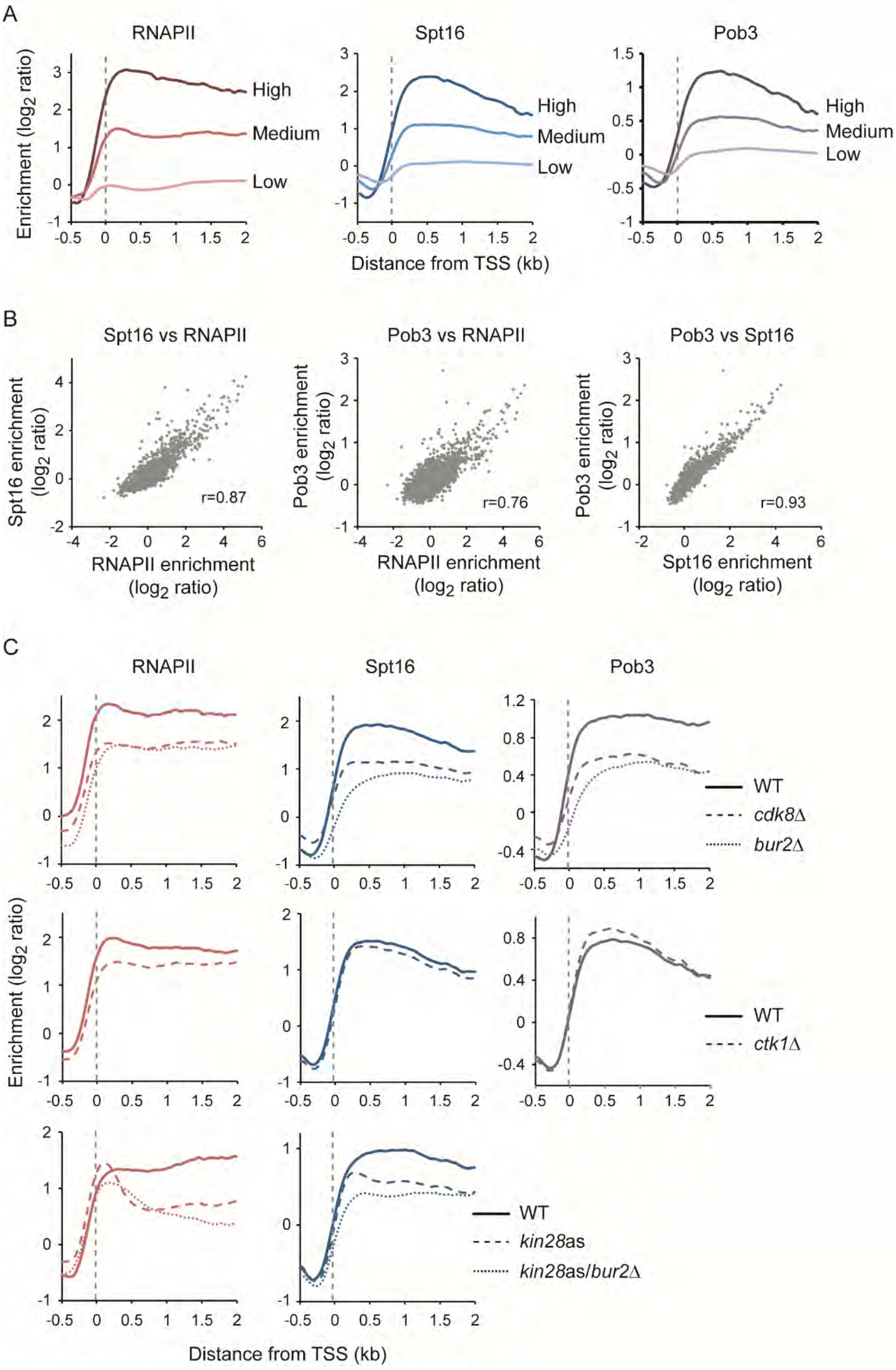
A
RNAPII
Spt16
Pob3
Enrichment (log2 ratio)
High
Medium
Low
Distance from TSS (kb)
B
Spt16 vs RNAPII
Pob3 vs RNAPII
Pob3 vs Spt16
Spt16 enrichment (log2 ratio)
Pob3 enrichment (log2 ratio)
RNAPII enrichment (log2 ratio)
Spt16 enrichment (log2 ratio)
r=0.87
r=0.76
r=0.93
C
RNAPII
Spt16
Pob3
WT
cdk8Δ
bur2Δ
WT
ctk1Δ
WT
kin28as
kin28as/bur2Δ
Enrichment (log2 ratio)
Distance from TSS (kb)

**Figure S1**: FACT occupancy correlates with transcription by RNAPII but that does not require CTD phosphorylation, related to **Figure 1**. **A**) Metagenes of RNAPII (Rpb3) and FACT (Spt16, Pob3) occupancy over highly (dark shade; n = 85), mildly (medium shade; n = 190) and lowly (light shade; n = 2,141) transcribed genes, as determined by ChIP-chip, relative to Input, in WT cells. **B**) Scatter plots of Spt16 versus RNAPII (Rpb3; left), Pob3 versus RNAPII (Rpb3; middle), and Pob3 versus Spt16 (right) occupancy on ORFs (n = 6,692), as determined by ChIP-chip, relative to Input, in WT cells. **C**) Metagenes of RNAPII (Rpb3) and FACT (Spt16, Pob3) occupancy over transcribed genes (n = 275), as determined by ChIP-chip, relative to Input. Top panels, WT (solid traces), *cdk8*Δ (dashed traces) and *bur2*Δ (dotted traces) cells; Middle panels, WT (solid traces) and *ctk1*Δ (dashed traces) cells; Bottom panels, WT (solid traces), *kin28*as (dashed traces) and *kin28*as/*bur2*Δ (dotted traces) cells. WT, *kin28*as, and *kin28*as/*bur2*Δ cells in the bottom panels were treated 15 min with NAPP1.

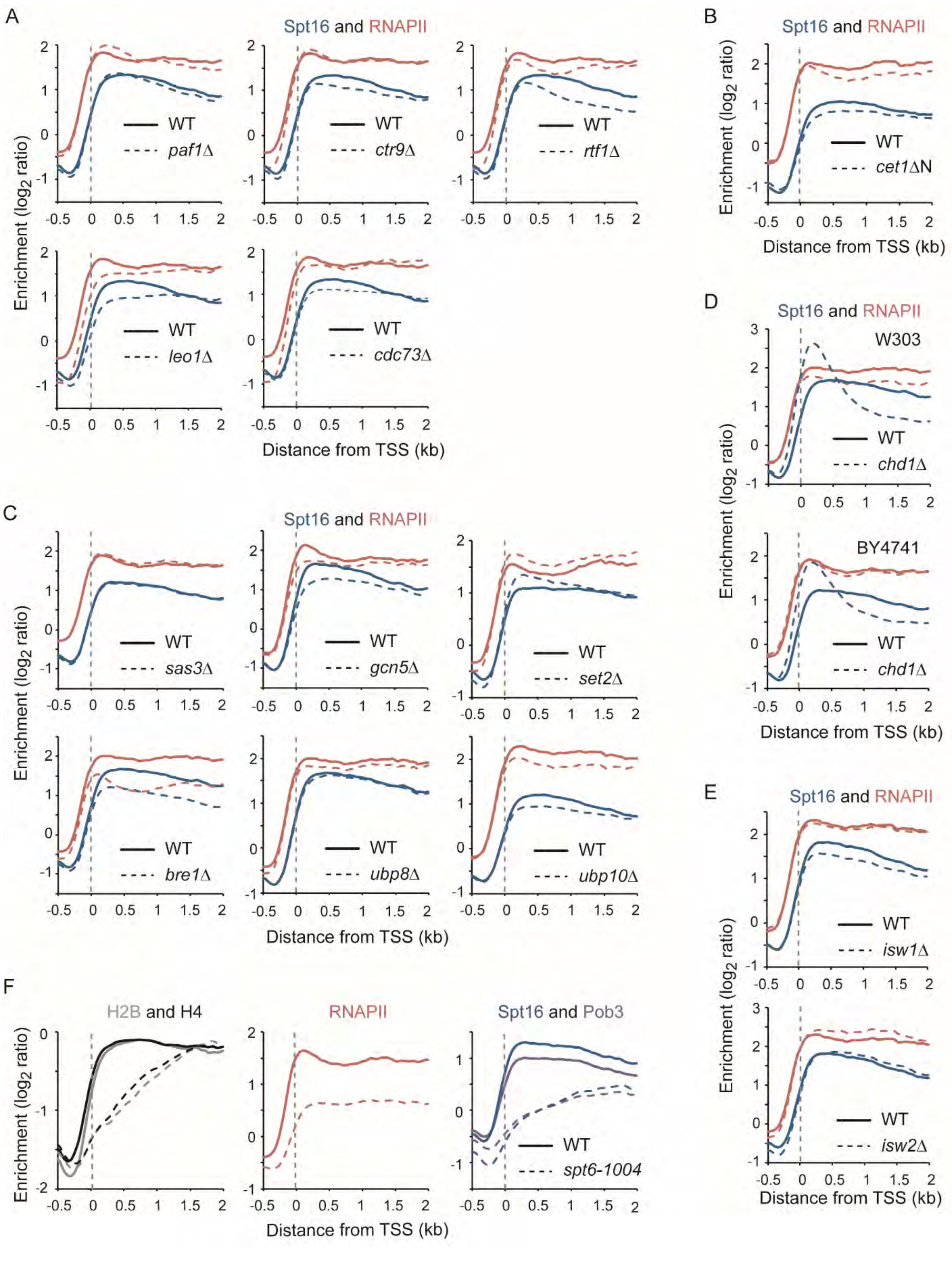

A
Spt16 and RNAPII
WT
paf1Δ
ctr9Δ
rtf1Δ
leo1Δ
cdc73Δ
Enrichment (log2 ratio)
Distance from TSS (kb)
B
Spt16 and RNAPII
WT
cet1ΔN
Enrichment (log2 ratio)
Distance from TSS (kb)
C
Spt16 and RNAPII
WT
sas3Δ
gcn5Δ
set2Δ
bre1Δ
ubp8Δ
ubp10Δ
Enrichment (log2 ratio)
Distance from TSS (kb)
D
Spt16 and RNAPII
W303
BY4741
WT
chd1Δ
Enrichment (log2 ratio)
Distance from TSS (kb)
E
Spt16 and RNAPII
WT
isw1Δ
isw2Δ
Enrichment (log2 ratio)
Distance from TSS (kb)
F
H2B and H4
RNAPII
Spt16 and Pob3
WT
spt6-1004
Enrichment (log2 ratio)
Distance from TSS (kb)

**Figure S2**: FACT recruitment is largely independent of PAF1C, the capping enzyme, chromatin modifiers, and chromatin remodelers but affected in *chd1*Δ cells, related to **Figure 2**. **A**) Metagenes of FACT (Spt16) and RNAPII (Rpb3) occupancy over transcribed genes (n = 275), as determined by ChIP-chip, relative to Input, in WT (solid traces) and PAF1C mutant (*paf1*Δ, *leo1*Δ, *ctr9*Δ, *cdc73*Δ, *rtf1*Δ; dashed traces) cells. **B**) Metagenes of FACT (Spt16) and RNAPII (Rpb3) occupancy over transcribed genes (n = 275), as determined by ChIP-chip, relative to Input, in WT (solid traces) and *cet1*ΔN (dashed traces) cells. **C**) Metagenes of FACT (Spt16) and RNAPII (Rpb3) occupancy over transcribed genes (n = 275), as determined by ChIP-chip, in WT (solid traces), sas3Δ, *gcn5*Δ, *set2*Δ, *bre1*Δ, *ubp8*Δ and *ubp10*Δ (dashed traces) cells. **D**) Metagenes of FACT (Spt16) and RNAPII (Rpb3) occupancy over transcribed genes (n = 275), as determined by ChIP-chip, relative to Input, in WT (solid traces) and *chd1*Δ (dashed traces) cells from W303 and BY4741 genetic backgrounds. **E**) Metagenes of FACT (Spt16) and RNAPII (Rpb3) occupancy over transcribed genes (n = 275), as determined by ChIP-chip, relative to Input, in WT (solid traces), *isw1*Δ (dashed traces, top panel) and *isw2*Δ (dashed traces, bottom panel) cells. **F**) Metagenes of histones H2B and H4, RNAPII (Rpb3) and FACT (Spt16, Pob3) occupancy over transcribed genes (n = 275), as determined by ChIP-chip, relative to Input, in WT (solid traces) and *spt6-1004* (dashed traces) cells which were shifted to 37°C for 80 min. Data for H2B and H4 are from (Jeronimo et al., 2019) and (Jeronimo et al., 2015), respectively.

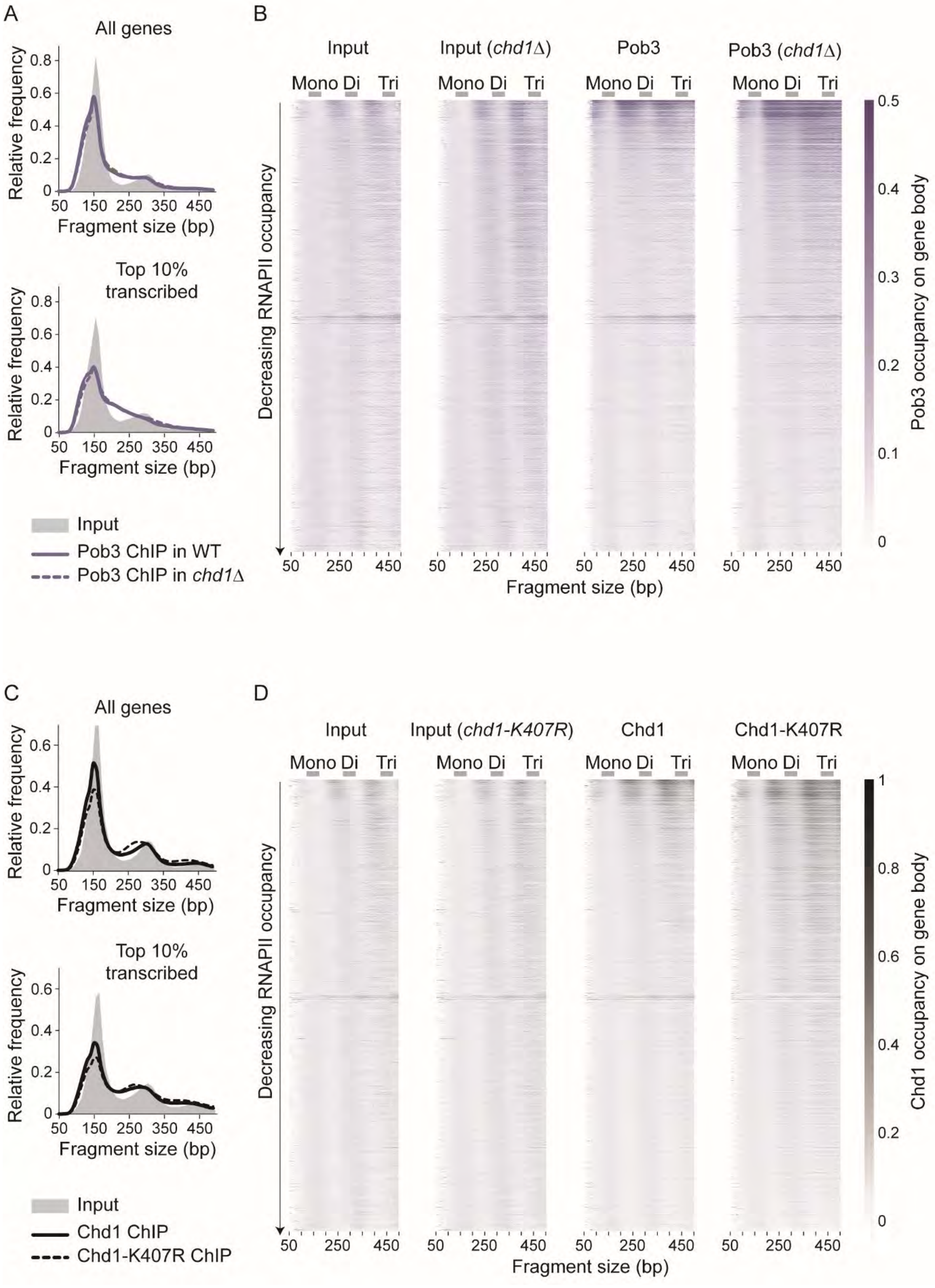

A
All genes
Relative frequency
0.8
0.6
0.4
0.2
0
50 150 250 350 450
Fragment size (bp)
Top 10% transcribed
Input
Pob3 ChIP in WT
Pob3 ChIP in chd1Δ
B
Input
Input (chd1Δ)
Pob3
Pob3 (chd1Δ)
Mono Di Tri
Decreasing RNAPII occupancy
Fragment size (bp)
Pob3 occupancy on gene body
C
All genes
Top 10% transcribed
Input
Chd1 ChIP
Chd1-K407R ChIP
D
Input
Input (chd1-K407R)
Chd1
Chd1-K407R
Chd1 occupancy on gene body

**Figure S3**: Analysis of the size of the mapped DNA fragments from the FACT (Pob3) and Chd1 MNase-ChIP-seq experiments, related to **Figure 3**. **A**) Distribution plots of the size of DNA fragments recovered from Input (grey area) and Pob3 (violet traces) MNase-ChIP-seq samples from WT (solid traces) and *chd1*Δ (dotted traces) cells. Top panel shows data for all genes (n = 5,796) and bottom panel shows data for the top 10% transcribed genes (n = 580) genes. **B**) Heatmap representation of Input and Pob3 average occupancy on gene body (n = 5,796, sorted by decreasing RNAPII occupancy on the y-axis) from WT and *chd1*Δ cells, as determined by MNase-ChIP-seq, computed with DNA fragments from different sizes (x-axis). **C**) Distribution plots of the size of DNA fragments recovered from Input (grey area) and Chd1 (black traces) MNase- ChIP-seq samples from WT (solid traces) and *chd1*-K407R (dotted traces) cells. Top panel shows data for all genes (n = 5,796) and bottom panel shows data for the top 10% transcribed genes (n = 580) genes. **D**) Heatmap representation of Input and Chd1 average occupancy on gene body (n = 5,796, sorted by decreasing RNAPII occupancy on the y-axis) from WT and *chd1*-K407R cells, as determined by MNase- ChIP-seq, computed with DNA fragments from different sizes (x-axis). Mono-, di- and tri-nucleosome-sized DNA fragments are indicated.

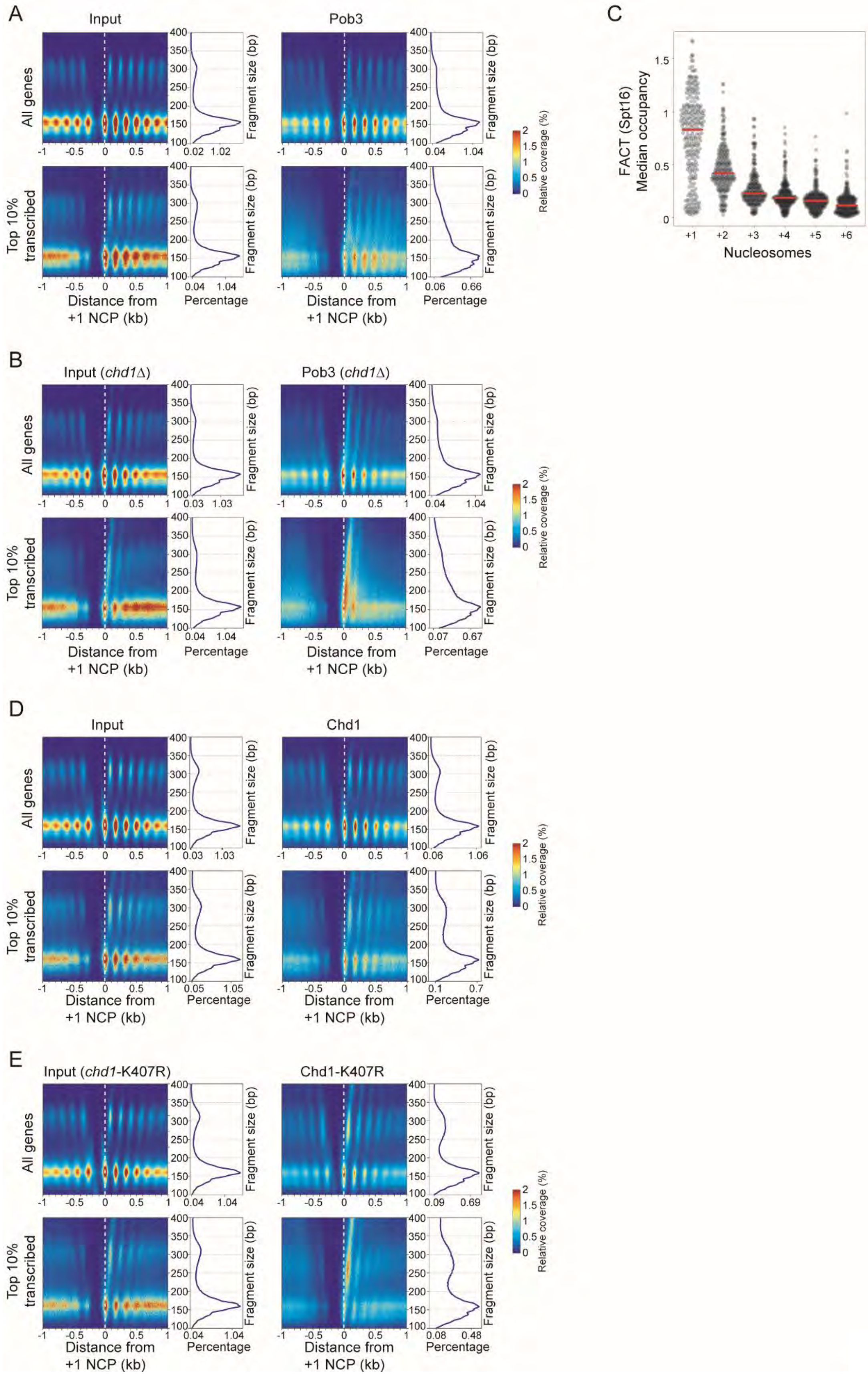
A
Input
Pob3
All genes
Top 10% transcribed
Fragment size (bp)
Distance from +1 NCP (kb)
Percentage
Relative coverage (%)
0.02 1.02
0.04 1.04
0.04 1.04
0.06 0.66
B
Input (chd1Δ)
Pob3 (chd1Δ)
0.03 1.03
0.04 1.04
0.04 1.04
0.07 0.67
C
FACT (Spt16) Median occupancy
Nucleosomes
+1 +2 +3 +4 +5 +6
D
Input
Chd1
0.03 1.03
0.06 1.06
0.05 1.05
0.1 0.7
E
Input (chd1-K407R)
Chd1-K407R
0.04 1.04
0.09 0.69
0.04 1.04
0.08 0.48

**Figure S4**: 2DO plots for Pob3 and Chd1 MNase-ChIP-seq experiments, related to **Figure 4**. **A**) 2DO plots of the coverage of the sequenced fragment mid-points (nucleosome dyads) in Input (left) and Pob3 ChIP (right) from MNase-digested chromatin from WT cells, relative to the +1 nucleosome dyads. On the right of each heatmap is the distribution of the fragment sizes. Top panels show data for all genes (n = 5,796) and bottom panels show data for the most transcribed genes (n = 580, top 10%). **B**) 2DO plots of the coverage of the sequenced fragment mid-points (nucleosome dyads) in Input (left) and Pob3 ChIP (right) from MNase-digested chromatin from *chd1*Δ cells, relative to the +1 nucleosome dyads. On the right of each heatmap is the distribution of the fragment sizes. Top panels show data for all genes (n = 5,796) and bottom panels show data for the most transcribed genes (n = 580, top 10%). **C**) Violin plot of the median signal from Spt16 MNase-ChIP-seq in *chd1*Δ cell (data shown in **Figure 4B**) over the six first nucleosomes for the top 10% transcribed genes (n = 580). The occupancies are calculated for all fragments in the 200-300 bp interval and using a nucleosome repeat length of 161 bp to delineate nucleosome territories. Red bars indicate the median. **D**) 2DO plots of the coverage of the sequenced fragment mid-points (nucleosome dyads) in Input (left) and Chd1 ChIP (right) from MNase-digested chromatin from WT cells, relative to the +1 nucleosome dyads. On the right of each heatmap is the distribution of the fragment sizes. Top panels show data for all genes (n = 5,796) and bottom panels show data for the most transcribed genes (n = 580, top 10%). **E**) 2DO plots of the coverage of the sequenced fragment mid-points (nucleosome dyads) in Input (left) and Chd1 ChIP (right) from MNase-digested chromatin from *chd1*-K407R cells, relative to the +1 nucleosome dyads. On the right of each heatmap is the distribution of the fragment sizes. Top panels show data for all genes (n = 5,796) and bottom panels show data for the most transcribed genes (n = 580, top 10%).

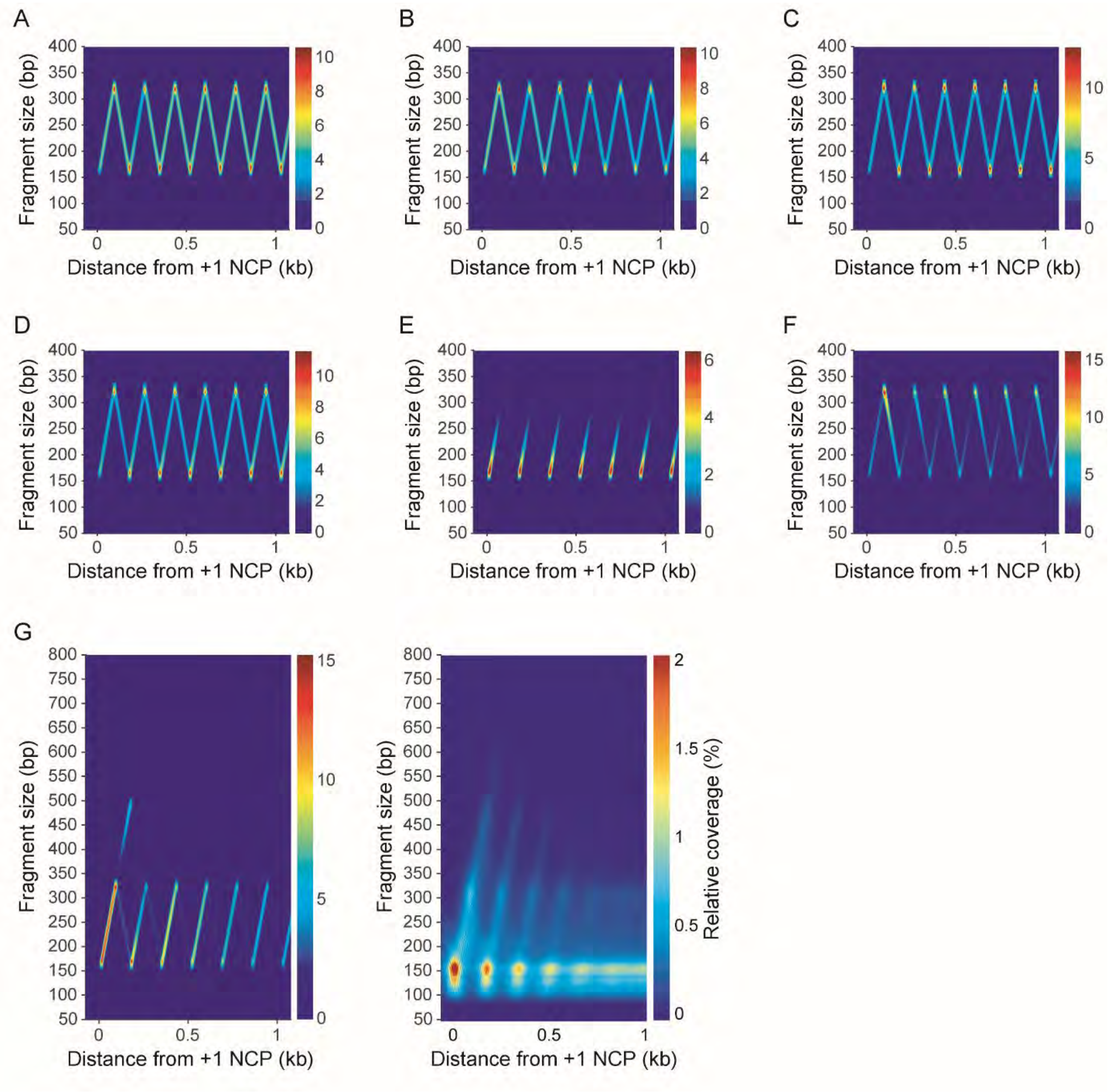


**Figure S5**: Mathematical modeling indicated that many plausible mechanisms are not a good fit to the observed experimental data, related to **Figure 5**. **A**) The most basic version of the model with FACT binding at the +1 nucleosome location, extension, and retraction. **B**) The basic version of the model, as in A), with constant FACT unbinding. **C**) The basic version of the model, as in A), with reduced rates of transition between the extending and retracting states. **D**) The full version of the model as in **Figure 5A** but with mid-gene loading and constant unbinding. **E**) The version of the model as in D) but with higher unbinding. **F**) The full version of the model as in **Figure 5A** but with mid-gene loading, constant unbinding, and a reduced retraction rate. **G**) Left: The full version of the model with reduced extension to match the *chd1*Δ mutant (as in **Figure 5D**) but depicting with larger fragment sizes up to 800 bp. Right: The experimental data for *chd1*Δ cells (same as in **Figure 4B**, top right panel) but depicting with larger fragment sizes up to 800 bp. The units of the color bars in the simulation panels represent the number of fragments observed in a complete simulation averaged over a 20x20 bp window (x$10^5$).

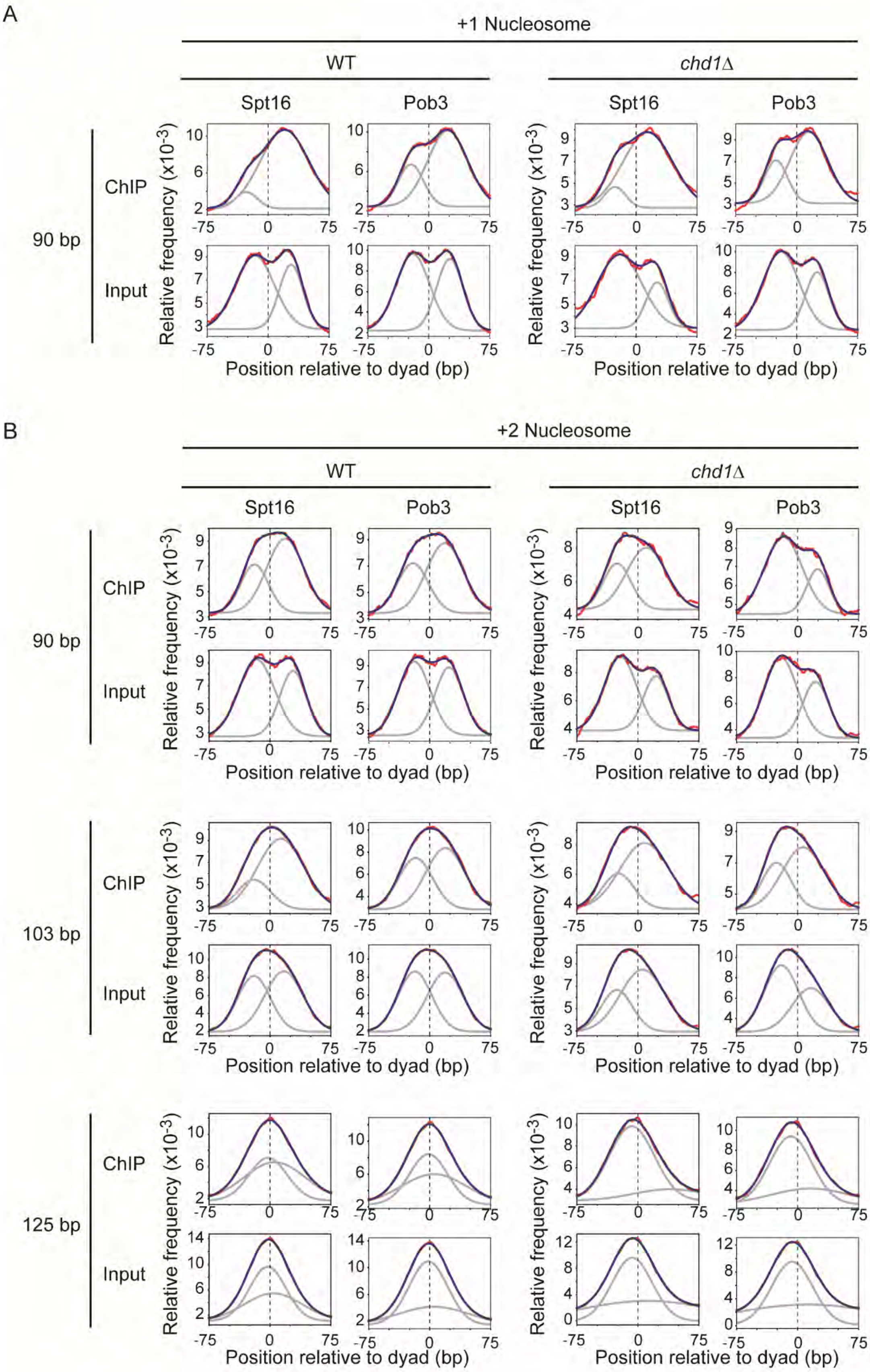
A
+1 Nucleosome
WT
chd1Δ
Spt16
Pob3
Spt16
Pob3
90 bp
ChIP
Input
Relative frequency (x10-3)
Position relative to dyad (bp)
B
+2 Nucleosome
WT
chd1Δ
Spt16
Pob3
Spt16
Pob3
90 bp
103 bp
125 bp
ChIP
Input
Relative frequency (x10-3)
Position relative to dyad (bp)
-75
0
75

**Figure S6:** Distribution of subnucleosomal-size DNA fragments from FACT MNase-ChIP-seq experiments, relative to +1 and +2 nucleosome dyads, related to **Figure 6**. **A**) Distribution of 90 bp (+/- 5 bp) fragment centers (red), from Spt16 and Pob3 MNase- ChIP-seq experiments (and their Inputs) from WT and *chd1*Δ cells, plotted relative to the position of +1 nucleosome dyads. The data (red) were fitted to a double-Gaussian (blue). The grey traces show the two individual Gaussians. **B**) Distribution of 90, 103, and 125 bp (+/- 5 bp) fragment centers (red), from Spt16 and Pob3 MNase-ChIP-seq experiments (and their Inputs) from WT and *chd1*Δ cells, plotted relative to the position of +2 nucleosome dyads. The data was fitted to a double-Gaussian (blue). The grey traces show the two individual Gaussians.

**Table S1. Yeast strains used in this study. Related to STAR Methods.**

| **Strain** | **Alias** | **Genotype** | **Source** |
|---|---|---|---|
| yFR116 | Z1256 | *MAT**a**, ade2-1, trp1-1, can1-100, leu2-3,112, his3-11,15, ura3* | (Ren et al., 2000) |
| yFR117 | BY4741 | *MAT**a**, his3Δ1, leu2Δ0, met15Δ0, ura3Δ0* | Open Biosystems |
| yFR1221 | DB1033 | *MATα, ura3-52, SUC2+* | (Nonet et al., 1987) |
| yFR1214 | Y260 | *MAT**a**, ura3-52, rpb1-1* | (Nonet et al., 1987) |
| yFR762 | SHY278 | *MATα, ade2::hisG, his3Δ200, leu2Δ0, lys2Δ0, met15Δ0, trp1Δ63, ura3Δ0* | (Liu et al., 2004) |
| yFR763 | SHY508 A | *MATα, ade2::hisG, his3Δ200, leu2Δ0, lys2Δ0, met15Δ0, trp1Δ63, ura3Δ0, kin28::kin28as(L83G) [pSH579, CEN, URA3, kin28as(L83G)]* | (Liu et al., 2004) |
| yFR1499 | | *MATα, ade2-1, trp1-1, can1-100, leu2-3,112, his3-11,15, ura3, tor1-1, fpr1Δ::NAT, RPL13A-2xFKBP12::TRP1, RPB1-FRB::KanMX6, [pFR482 (CEN, HIS3, RPB1-CTDWT-3xFLAG)]* | (Jeronimo and Robert, 2014) |
| yFR1501 | | *MATα, ade2-1, trp1-1, can1-100, leu2-3,112, his3-11,15, ura3, tor1-1, fpr1Δ::NAT, RPL13A-2xFKBP12::TRP1, RPB1-FRB::KanMX6, [pFR486 (CEN, HIS3, rpb1-CTDS2A-3xFLAG)]* | (Jeronimo and Robert, 2014) |
| yFR1503 | | *MATα, ade2-1, trp1-1, can1-100, leu2-3,112, his3-11,15, ura3, tor1-1, fpr1Δ::NAT, RPL13A-2xFKBP12::TRP1, RPB1-FRB::KanMX6, [pFR490 (CEN, HIS3, rpb1-CTDS5A-3xFLAG)]* | (Jeronimo and Robert, 2014) |
| yFR1526 | | *MATα, ade2-1, trp1-1, can1-100, leu2-3,112, his3-11,15, ura3, tor1-1, fpr1Δ::NAT, RPL13A-2xFKBP12::TRP1, RPB1-FRB::KanMX6, [pFR494 (CEN, HIS3, rpb1-CTDS7A-3xFLAG)]* | (Jeronimo and Robert, 2014) |
| yFR999 | | *MATa, ade2-1, trp1-1, can1-100, leu2-3,112, his3-11,15, ura3, cdk8Δ::URA3* | (Bataille et al., 2012) |
| yFR682 | | *MATa, his3Δ1, leu2Δ0, met15Δ0, ura3Δ0, bur2Δ::kanMX4* | (Bataille et al., 2012) |
| yFR551 | | *MATa, his3Δ1, leu2Δ0, met15Δ0, ura3Δ0, ctk1Δ::kanMX4* | (Bataille et al., 2012) |

| | | | |
|---|---|---|---|
| yFR912 | | *MATα, ade2::hisG, his3Δ200, leu2Δ0, lys2Δ0, met15Δ0, trp1Δ63, ura3Δ0, bur2Δ::LEU2, kin28::kin28as(L83G) [pSH579, CEN, URA3, kin28as(L83G)]* | (Bataille et al., 2012) |
| yFR930 | | *MAT**a**, ade2-1, trp1-1, can1-100, leu2-3,112, his3-11,15, ura3, chd1Δ::URA3* | (Jeronimo et al., 2019) |
| yFR3061 | | *MAT**a**, ade2-1, trp1-1, can1-100, leu2-3,112, his3-11,15, ura3, CHD1::3HA-CHD1* | This study |
| yFR3084 | | *MAT**a**, ade2-1, trp1-1, can1-100, leu2-3,112, his3-11,15, ura3, CHD1::chd1-K407R* | This study |
| yFR3075 | | *MAT**a**, ade2-1, trp1-1, can1-100, leu2-3,112, his3-11,15, ura3, CHD1::3HA-chd1-K407R* | This study |
| yFR1350 | | *MAT**a**, his3Δ1, leu2Δ0, met15Δ0, ura3Δ0, paf1Δ::KanMX4* | Open Biosystems |
| yFR1368 | | *MAT**a**, his3Δ1, leu2Δ0, met15Δ0, ura3Δ0, ctr9Δ::KanMX4* | Open Biosystems |
| yFR1552 | | *MAT**a**, his3Δ1, leu2Δ0, met15Δ0, ura3Δ0, leo1Δ::KanMX4* | Open Biosystems |
| yFR3208 | | *MAT**a**, his3Δ1, leu2Δ0, met15Δ0, ura3Δ0, cdc73Δ::URA3* | This study |
| yFR3209 | | *MAT**a**, his3Δ1, leu2Δ0, met15Δ0, ura3Δ0, rtf1Δ::URA3* | This study |
| yFR3215 | YSB709 | *MAT**a**, ura3-52, leu2Δ1, trp1Δ63, his3Δ200, lys2Δ202, cet1Δ1::TRP1 [pRS315-CET1]* | (Takase et al., 2000) |
| yFR3216 | YSB710 | *MATa, ura3-52, leu2Δ1, trp1Δ63, his3Δ200, lys2Δ202, cet1Δ1::TRP1 [pRS315-CET1(Pro+205-549)]* | (Takase et al., 2000) |
| yFR297 | | *MATa, his3Δ1, leu2Δ0, met15Δ0, ura3Δ0, chd1Δ::KanMX4* | Open Biosystems |
| yFR729 | | *MAT**a**, ade2-1, trp1-1, can1-100, leu2-3,112, his3-11,15, ura3, isw1Δ::URA3* | This study |
| yFR928 | | *MAT**a**, ade2-1, trp1-1, can1-100, leu2-3,112, his3-11,15, ura3, isw2Δ::LEU2* | This study |
| yFR972 | FY80 | *MAT**a**, leu2Δ1, lys2-128δ, trp1Δ 63, ura3-52* | (Cheung et al., 2008) |
| yFR974 | FY2425 | *MATα, his3Δ200, leu2Δ1, lys2-128δ, ura3-52, spt6-1004-FLAG* | (Cheung et al., 2008) |
| yFR083 | | *MATa, his3Δ1, leu2Δ0, met15Δ0, ura3Δ0, sas3ΔkanMX4* | R.A. Young |
| yFR265 | | *MATa, his3Δ1, leu2Δ0, met15Δ0, ura3Δ0, gcn5ΔkanMX4* | Open Biosystems |
| yFR348 | | *MAT**a**, ade2-1, trp1-1, can1-100, leu2-3,112, his3-11,15, ura3, set2Δ::HIS3* | (Drouin et al., 2010) |
| yFR2024 | | *MATa, ade2-1, trp1-1, can1-100, leu2-3,112, his3-11,15, ura3, bre1Δ::LEU2* | This study |

| | | | |
|---|---|---|---|
| yFR2035 | | *MATa, ade2-1, trp1-1, can1-100, leu2-3,112, his3-11,15, ura3, ubp8D::KanMX4* | This study |
| yFR3198 | | *MAT**a**, ade2-1, trp1-1, can1-100, leu2-3,112, his3-11,15, ura3, ubp10Δ::KanMX4* | This study |
| yFR705 | | *MAT**a**, ade2-1, trp1-1, can1-100, leu2-3,112, his3-11,15, ura3, spt4Δ::URA3* | (Drouin et al., 2010) |
| yFR721 | | *MAT**a**, his3Δ1, leu2Δ0, met15Δ0, ura3Δ0, spt4Δ::KanMX4* | Open Biosystems |

### Table S2. Oligonucleotides used in this study. Related to STAR Methods.

| **Description** | **Sequence** | **Source** |
|---|---|---|
| **CRISPR oligos** | | |
| Oligos used to clone gDNA targetting *CHD1* in pML107 (to introduce the K407R mutation) | Foward: GATCAACTGATAAAGGCGACAGTCGTTTTAGAGCTAG<br><br>Reverse: CTAGCTCTAAAACGACTGTCGCCTTTATCAGTT | This study |
| Oligos used to generate a donor DNA introducing the K407R mutation in *CHD1* | Foward: GATGGCATTTTTGTGGTCCAAAGGTGATAATGGTATACTGGCAGATGAGATGGGCCTGGGACGTACGGTG<br><br>Reverse: GTCCGTTTTGTCTACGAGCAAATATCAGCCAACTGATAAAGGCGACAGTCTGCACCGTACGTCCCAGGCC | This study |
| **ChIP-chip oligos** | | |
| Oligos used to generate annealed linkers | Foward: GCGGTGACCCGGGAGATCTGAATTC<br><br>Reverse: GAATTCAGATC | (Ren et al., 2000) |
| **ChIP-exo oligos** | | |
| 1st ligation adapter (D701) | Exa2B: GAT CGG AAG AGC ACA CGT CTG AAC TCC AGT CAC<br><br>ExB2-i7_D701: /5Phos/CAA GCA GAA GAC GGC ATA CGA GAT ATTACTCG GTG ACT GGA GTT CAG ACG TGT GCT CTT CCG ATC T | This study |
| 1st ligation adapter (D702) | Exa2B: GAT CGG AAG AGC ACA CGT CTG AAC TCC AGT CAC<br><br>ExB2-i7_D702: /5Phos/CAA GCA GAA GAC GGC ATA CGA GAT TCCGGAGA GTG ACT GGA GTT CAG ACG TGT GCT CTT CCG ATC T | This study |
| 1st ligation adapter (D703) | Exa2B: GAT CGG AAG AGC ACA CGT CTG AAC TCC AGT CAC<br><br>ExB2-i7_D703: /5Phos/CAA GCA GAA GAC GGC ATA CGA GAT CGCTCATT GTG ACT GGA GTT CAG ACG TGT GCT CTT CCG ATC T | This study |

| | | |
|---|---|---|
| 1st ligation adapter (D704) | Exa2B: GAT CGG AAG AGC ACA CGT CTG AAC TCC AGT CAC<br><br>ExB2-i7_D704: /5Phos/CAA GCA GAA GAC GGC ATA CGA GAT GAGATTCC GTG ACT GGA GTT CAG ACG TGT GCT CTT CCG ATC T | This study |
| 2nd ligation adapter (D501) | ExA1-SSL_N5: NNN NNA GAT CGG AAG AGC G<br><br>ExC1-i5_D501: AAT GAT ACG GCG ACC ACC GAG ATC TAC AC TATAGCCT AC ACT CTT TCC CTA CAC GAC GCT CTT CCG ATC T | This study |
| 2nd ligation adapter (D502) | ExA1-SSL_N5: NNN NNA GAT CGG AAG AGC G<br><br>ExC1-i5_D502: AAT GAT ACG GCG ACC ACC GAG ATC TAC AC ATAGAGGC AC ACT CTT TCC CTA CAC GAC GCT CTT CCG ATC T | This study |
| 2nd ligation adapter (D503) | ExA1-SSL_N5: NNN NNA GAT CGG AAG AGC G<br><br>ExC1-i5_D503: AAT GAT ACG GCG ACC ACC GAG ATC TAC AC CCTATCCT AC ACT CTT TCC CTA CAC GAC GCT CTT CCG ATC T | This study |
| 2nd ligation adapter (D504) | ExA1-SSL_N5: NNN NNA GAT CGG AAG AGC G<br><br>ExC1-i5_D504: AAT GAT ACG GCG ACC ACC GAG ATC TAC AC GGCTCTGA AC ACT CTT TCC CTA CAC GAC GCT CTT CCG ATC T | This study |
| Oligos used to amplify ChIP-exo libraries | P1.3: AAT GAT ACG GCG ACC ACC<br><br>P2.1: CAA GCA GAA GAC GGC ATA CGA G | (Rossi et al., 2018) |

**Movie S1**: Animation of the "inchworm-like" model of FACT recruitment spreading for the WT case. The left panel shows a representation of the regions of DNA protected from MNase digestion by FACT complexes for 100 realisations of a simulation across a synthetic gene. White bars represent FACT complexes, the x-axis is location in the gene and each track on the y-axis represents an individual realisation (tracks are separated by blank spaces). The right panel shows the density of fragments, of given sizes (y-axis) and locations (x-axis), protected from MNase digestion by FACT complexes as they are sampled from the simulation. Parameters of the simulation are as for the WT case as in the main text (**Figure 5B** and **Table 1**) except that the number of time steps shown is $10^5$, there are 100 realisations and there is no relaxation period prior to measurements for the protected fragments being taken.

**Movie S2**: Animation of the "inchworm-like" model of FACT recruitment spreading for the *chd1*Δ case (same as WT except for a reduced extension rate of the FACT complex). The left panel shows a representation of the regions of DNA protected from MNase digestion by FACT complexes for 100 realisations of a simulation across a synthetic gene. White bars represent FACT complexes, the x-axis is location in the gene and each track on the y-axis represents an individual realisation (tracks are separated by blank spaces). The right panel shows the density of fragments, of given sizes (y-axis) and locations (x-axis), protected from MNase digestion by FACT complexes as they are sampled from the simulation. Parameters of the simulation are as for the *chd1*Δ case as in the main text (**Figure 5D** and **Table 1**) except that the number of time steps shown is $10^5$, there are 100 realisations and there is no relaxation period prior to measurements for the protected fragments being taken.

### REFERENCES


Bataille, A.R., Jeronimo, C., Jacques, P.E., Laramee, L., Fortin, M.E., Forest, A., Bergeron, M., Hanes, S.D., and Robert, F. (2012). A universal RNA polymerase II CTD cycle is orchestrated by complex interplays between kinase, phosphatase, and isomerase enzymes along genes. Mol Cell *45*, 158-170.

Cheung, V., Chua, G., Batada, N.N., Landry, C.R., Michnick, S.W., Hughes, T.R., and Winston, F. (2008). Chromatin- and transcription-related factors repress transcription from within coding regions throughout the Saccharomyces cerevisiae genome. PLoS Biol *6*, e277.

Drouin, S., Laramee, L., Jacques, P.E., Forest, A., Bergeron, M., and Robert, F. (2010). DSIF and RNA polymerase II CTD phosphorylation coordinate the recruitment of Rpd3S to actively transcribed genes. PLoS Genet *6*, e1001173.

Jeronimo, C., Poitras, C., and Robert, F. (2019). Histone Recycling by FACT and Spt6 during Transcription Prevents the Scrambling of Histone Modifications. Cell Rep *28*, 1206-1218 e1208.

Jeronimo, C., and Robert, F. (2014). Kin28 regulates the transient association of Mediator with core promoters. Nat Struct Mol Biol *21*, 449-455.

Jeronimo, C., Watanabe, S., Kaplan, C.D., Peterson, C.L., and Robert, F. (2015). The Histone Chaperones FACT and Spt6 Restrict H2A.Z from Intragenic Locations. Mol Cell *58*, 1113-1123.

Liu, Y., Kung, C., Fishburn, J., Ansari, A.Z., Shokat, K.M., and Hahn, S. (2004). Two cyclin-dependent kinases promote RNA polymerase II transcription and formation of the scaffold complex. Mol Cell Biol *24*, 1721-1735.

Nonet, M., Scafe, C., Sexton, J., and Young, R. (1987). Eucaryotic RNA polymerase conditional mutant that rapidly ceases mRNA synthesis. Mol Cell Biol *7*, 1602-1611.

Ren, B., Robert, F., Wyrick, J.J., Aparicio, O., Jennings, E.G., Simon, I., Zeitlinger, J., Schreiber, J., Hannett, N., Kanin, E., *et al.* (2000). Genome-wide location and function of DNA binding proteins. Science *290*, 2306-2309.

Rossi, M.J., Lai, W.K.M., and Pugh, B.F. (2018). Simplified ChIP-exo assays. Nat Commun *9*, 2842.

Takase, Y., Takagi, T., Komarnitsky, P.B., and Buratowski, S. (2000). The essential interaction between yeast mRNA capping enzyme subunits is not required for triphosphatase function in vivo. Mol Cell Biol *20*, 9307-9316.
